# Long-range coupled motions underlie ligand recognition by a chemokine receptor

**DOI:** 10.1101/2020.07.28.225664

**Authors:** Krishna Mohan Sepuru, Vinay Nair, Priyanka Prakash, Alemayehu A. Gorfe, Krishna Rajarathnam

**Affiliations:** Department of Biochemistry and Molecular Biology, The University of Texas Medical Branch, Galveston, Texas 77555; Department of Microbiology and Immunology, The University of Texas Medical Branch, Galveston, Texas 77555; Sealy Center for Structural Biology and Molecular Biophysics, The University of Texas Medical Branch, Galveston, Texas 77555; Integrative Biology and Pharmacology, McGovern Medical School, University of Texas Health Science Center at Houston, Houston, TX; MD Anderson Cancer Center UTHealth Graduate School of Biomedical Sciences, Houston, TX

**Keywords:** chemokine, CXCL8, GPCR, NMR, solution structure, Molecular Dynamics, long-range coupling, CXCR1

## Abstract

Chemokines are unusual class-A GPCR agonists because of their large size (∼10 kDa) and binding at two distinct receptor sites: N-terminal domain (Site-I, unique to chemokines) and a groove defined by extracellular loop/transmembrane helices (Site-II, shared with all small molecule class-A ligands). Whereas binding at Site-II triggers receptor activation, the role of Site-I is not known. Structures and sequence analysis reveal that the receptor N-terminal domains (N-domains) are flexible and contain intrinsic disorder. Using a hybrid NMR-MD approach, we characterized the role of Site-I interactions for the CXCL8-CXCR1 pair. NMR data indicate that the CXCR1 N-domain becomes structured on binding and that the binding interface is extensive with 30% of CXCL8 residues participating in this initial interaction. MD simulations indicate that CXCL8 bound at Site-I undergoes extensive reorganization on engaging Site-II with several residues initially engaged at Site-I also engaging Site-II. We conclude that structural plasticity of Site-I interactions plays an active role in driving ligand recognition by a chemokine receptor.

---

Chemokines, the largest subfamily of cytokines, play diverse and fundamental roles, from trafficking immune cells and organogenesis to combating infection and injury. Not surprisingly, a dysregulation in chemokine function has been implicated in the pathophysiology of various autoimmune and inflammatory diseases, and in cancer (Griffith et al., 2014; Nagarsheth et al., 2017). Chemokines mediate their function by activating receptors that belong to the G protein-coupled receptor (GPCR) class. Humans express around 50 chemokines and 20 receptors, which show a rich tapestry of interactions resulting in large differences in affinity, specificity, and activity. Some receptors are activated by multiple chemokines, and at the other end of the spectrum, some are strictly activated by a single chemokine. Chemokines, in turn, show a range of receptor specificity and activity. Diversity of interactions is further compounded by chemokines and chemokine receptors existing as monomers and dimers, biased G protein vs. β-arrestin signaling, interacting with glycosaminoglycans, post-translational modification, and binding to atypical chemokine receptors (Bhusal et al., 2020; Rajagopal et al., 2013; Rajarathnam and Desai, 2020).

At a structural level, within this bewildering complexity is also an elegant simplicity. Chemokines, despite large differences in sequence identity, share the same structural fold. The chemokine structure consists of a short stretch of N-terminal residues preceding the conserved cysteines, an N-terminal loop (N-loop) followed by three β-strands and an α-helix, and disulfides that link the N-terminal and N-loop residues to the structural core. As class A GPCRs, chemokine receptor structure consists of seven transmembrane (TM) helices, three extracellular (EC) and three intracellular (IC) loops, and a short extracellular N-terminal and an intracellular C-terminal domain. Therefore, the spectrum of functional phenotypes must be encoded at the amino acid sequence level on a conserved ligand and receptor structural scaffolds.

Chemokines are unusual agonists for class A GPCRs, as conventional agonists tend to be small molecules with a rigid scaffold such as neurotransmitters, hormones, and those related to smell and taste. Chemokines are much bigger (MW ∼10 kDa) but functionally no different as they bind their receptors with nanomolar affinity and activate the same signaling pathways. Therefore, chemokines must have evolved to exploit the GPCR structural features. Chemokines bind the receptor at two distinct sites: the N-terminal domain (Site-I, unique to chemokines) and a groove defined by the receptor EC loops/TM helices (Site-II, shared with all class A ligands) (Crump et al., 1997; Rajagopalan and Rajarathnam, 2006; Thiele and Rosenkilde, 2014). Whereas chemokine binding at Site-II triggers receptor activation, the role of Site-I interactions is not known.

In this study, we provide a structural model of how Site-I interactions orchestrate CXCL8 monomer binding to the CXCR1 receptor. Chemokines are classified into CXC, CC, CX_3_C, and C subfamilies on the basis of conserved cysteines in the N-terminus. CXCL8, as a member of a subset of seven CXC chemokines characterized by the N-terminal ‘ELR’ motif, activates CXCR1 and CXCR2 receptors. Whereas all ELR-chemokines are potent CXCR2 agonists, CXCL8 monomer alone functions as a potent CXCR1 agonist (Nasser et al., 2009). CXCR1 and CXCR2 are expressed in neutrophils and in non-immune cells such as hepatocytes and neuronal cells, and these receptors play fundamental roles in combating infection, regulating pain, and in the trafficking and proliferation of cancer cells (Ha et al., 2017; Rajarathnam et al., 2019). CXCR1 and CXCR2 sequences show large differences for the N-domain indicating a prominent role for Site-I interactions in determining specificity and affinity.

Knowledge of the structures are essential to describe the molecular mechanisms by which Site-I interactions determine binding at Site-II. The Site-I N-domain residues are not observed in the CXCR1 structure (Park et al., 2012). Sequences predict intrinsic disorder is a shared feature of N-domain of not only chemokine receptors but all class A GPCRs (Latorraca et al., 2017; Raucci et al., 2014; Venkatakrishnan et al., 2014). Characteristic features of intrinsic disorder include a preponderance of polar and charged residues, lack of a hydrophobic core, engaging in transient but specific interactions, and becoming structured on binding a protein partner (Latysheva et al., 2015; Uversky, 2019; van der Lee et al., 2014; Wright and Dyson, 2015). The N-domain residues are also not observed in the crystal structures of inhibitor-bound chemokine receptor complexes (Liu et al., 2020; Wu et al., 2010). The structure of CXCL8-CXCR1 complex is not known, but cryoelectron microscopy (cryoEM) and crystal structures of CXCL8 bound to CXCR2 and also of chemokines bound to CXCR4, CCR5, and CCR6 receptors are known (Liu et al., 2020; Qin et al., 2015; Wasilko et al., 2020; Zheng et al., 2017). A significant segment of the receptor N-domain is also missing in these structures, suggesting Site-I interactions are either weak, lost once the chemokine engages Site-II residues, or that the receptor Site-I residues continue to be dynamic in the chemokine-bound form.

We propose a hybrid strategy that combines nuclear magnetic resonance (NMR) spectroscopy for determining the solution structure of the CXCL8 monomer-CXCR1 N-domain complex and extended molecular dynamics (MD) simulations to describe how CXCL8 bound at Site-I engages Site-II residues. Our model for Site-I interactions is as follows. The first step involves CXCR1 N-domain capturing CXCL8 by the fly-casting mechanism (Shoemaker et al., 2000), as the unstructured N-domain has a large capture radius. CXCL8 bound at Site-I is a distinct but transient intermediate, and in the next step, Site-I functions as an anchor for CXCL8 N-terminal residues to search for Site-II residues. CXCL8 binding the CXCR1 N-domain captures the structural features of the Site-I intermediate. The NMR structure of the Site-I complex reveals an extensive binding interface stabilized by packing and ionic interactions. Interestingly, MD simulations reveal that Site-II interactions are coupled to extensive reorganization of Site-I interactions, including disengagement, rearrangement, and several residues initially engaged in Site-I interactions alone engaging in both Site-I and Site-II interactions. Our data make a compelling case for long-range coupled motions in driving ligand recognition by the CXCR1 receptor.

## RESULTS

### Structure of the CXCL8-CXCR1 N-domain complex

The solution structure of the CXCL8 monomer bound to the CXCR1 N-domain Site-I complex was determined using 1310 distance and 106 angular restraints. Ensemble structures, structural details, and statistics are shown in Fig. 1, Supplementary Fig. 1, and Supplementary Tables 1 and 2. For the CXCL8 monomer, we used the CXCL8 variant (residues 1-66), which is missing the last six C-terminal residues known to stabilize the dimer interface (Berkamp et al., 2017; Joseph and Rajarathnam, 2015; Joseph et al., 2018). For CXCR1 N-domain, we used a construct containing residues 1 to 29 (Joseph and Rajarathnam, 2015; Joseph et al., 2018). To distinguish CXCR1 and CXCL8 residues, receptor residues are annotated as a single letter amino acid and ligand residues are annotated as a three-letter amino acid. The interface is extensive with as many as 30% of the CXCL8 residues (20 out of 66) participating in the Site-I interaction. The structure reveals that CXCL8 is more structured in the bound form. The backbone r.m.s.d. (root mean square deviation) of CXCL8 in the bound and free forms, for residues Lys11 to Leu66, is 0.34 and 0.65Å, respectively. The side chains of the interface residues highlight the overall higher conformational flexibility of CXCL8 in the free compared to the bound form (Figs. 1C, 1D). The CXCR1 N-domain peptide is unstructured in the free form. In the bound form, residues D11 to P29 are structured (r.m.s.d. 0.63Å), which is supported by 16 long-range intramolecular NOEs and 90 intermolecular NOEs to the CXCL8 N-loop, 3^rd^ β-strand (β_3_), and C-terminal helical residues (Figs. 1E, 1F). A schematic of CXCL8 and CXCR1 residues that show intermolecular NOEs is shown in Fig. 1G. The first ten distal residues of the CXCR1 N-domain are unstructured, which is evident from only a few sequential and no intermolecular NOEs.

**Figure 1.**
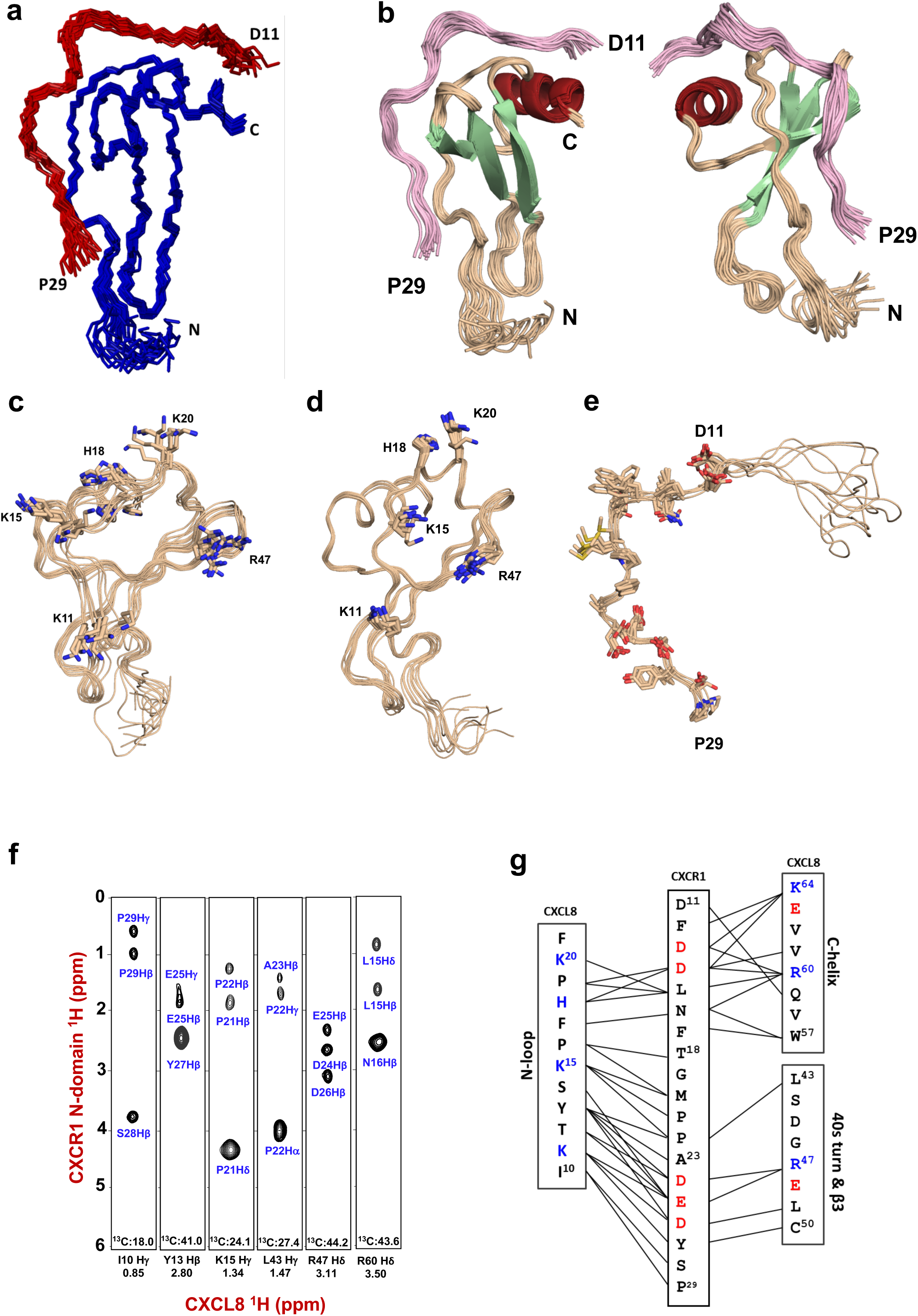
Solution structure of the CXCL8-CXCR1 N-domain complex. (**a)** Twenty ensemble backbone C_α_ traces of CXCL8-CXCR1 N-domain structure. For the CXCR1 N-domain, only the structured residues (D11 to P29) are shown. CXCL8 and CXCR1 N-domain are shown in blue and red, respectively. **(b)** An ensemble of the CXCL8-CXCR1 N-domain structures is shown in two different orientations for highlighting the binding interface. An ensemble of the CXCL8 monomer structure in the free (**c**) and CXCR1-bound (**d**) forms. The sidechains of a subset of basic residues that mediate Site-I interactions are shown and labeled. The nitrogens of the side chain are shown in blue. The ensemble indicates that the binding interface residues are structured in the bound form and conformationally more dynamic in the free form. (**e**) An ensemble of CXCR1 N-domain structure in the bound form with the side chains shown as a stick. Side chain nitrogens and oxygens are shown in blue and red, respectively. The first ten residues are unstructured and not involved in CXCL8 binding. (**f**) ^13^C-filter NOE strips representing intermolecular NOEs from CXCL8 N-loop, β_3_-strand, and α-helical residues to CXCR1 N-domain. (**g**) A schematic showing intermolecular NOE contacts between CXCR1 and CXCL8 N-loop, 40s turn/β_3_ and C-helix. Basic and acidic residues are in blue and red, respectively.

The structure reveals that the CXCR1 N-domain binds in an extended fashion with no defined secondary structure except for a type IV β-turn for residues F17 to M20. Extensive packing and 20 backbone-side chain and side chain-side chain H-bonds and salt bridges stabilize the complex. CXCR1 F12, F17, and P29 residues bind a hydrophobic groove defined by CXCL8 Tyr13, Phe17, Phe21, Trp57 and Val61 residues (Fig. 2A). This hydrophobic pocket is flanked by two ionic clusters (Figs. 2B, 2C). The first ionic cluster consists of a network of interactions between D11/Gln59, D13/Arg60, D14/Lys64, and D14/His18, and the second ionic cluster between D24/Arg47, E25/Lys11, and D26/Arg47. Interestingly, Lys15, which is unique to CXCL8 among ELR-chemokines, is not involved in ionic interactions, but is observed in the proximity of CXCR1 non-polar M20, P21, and P22 residues (Fig. 2D). NMR backbone ^15^N-relaxation measurements of the CXCL8-CXCR1 N-domain complex indicate that the CXCL8 N-loop residues undergo significant conformational fluctuations, CXCL8 N-terminal ELR residues are unstructured both in the free and bound forms, distal CXCR1 N-terminal residues 1 to 10 are unstructured, and CXCR1 residues 11 to 27 are structured but show significant dynamics (Joseph et al., 2018). These structural and dynamics characteristics collectively indicate that conformational flexibility, especially of the interface residues, promote Site-I complex formation.

**Figure 2.**
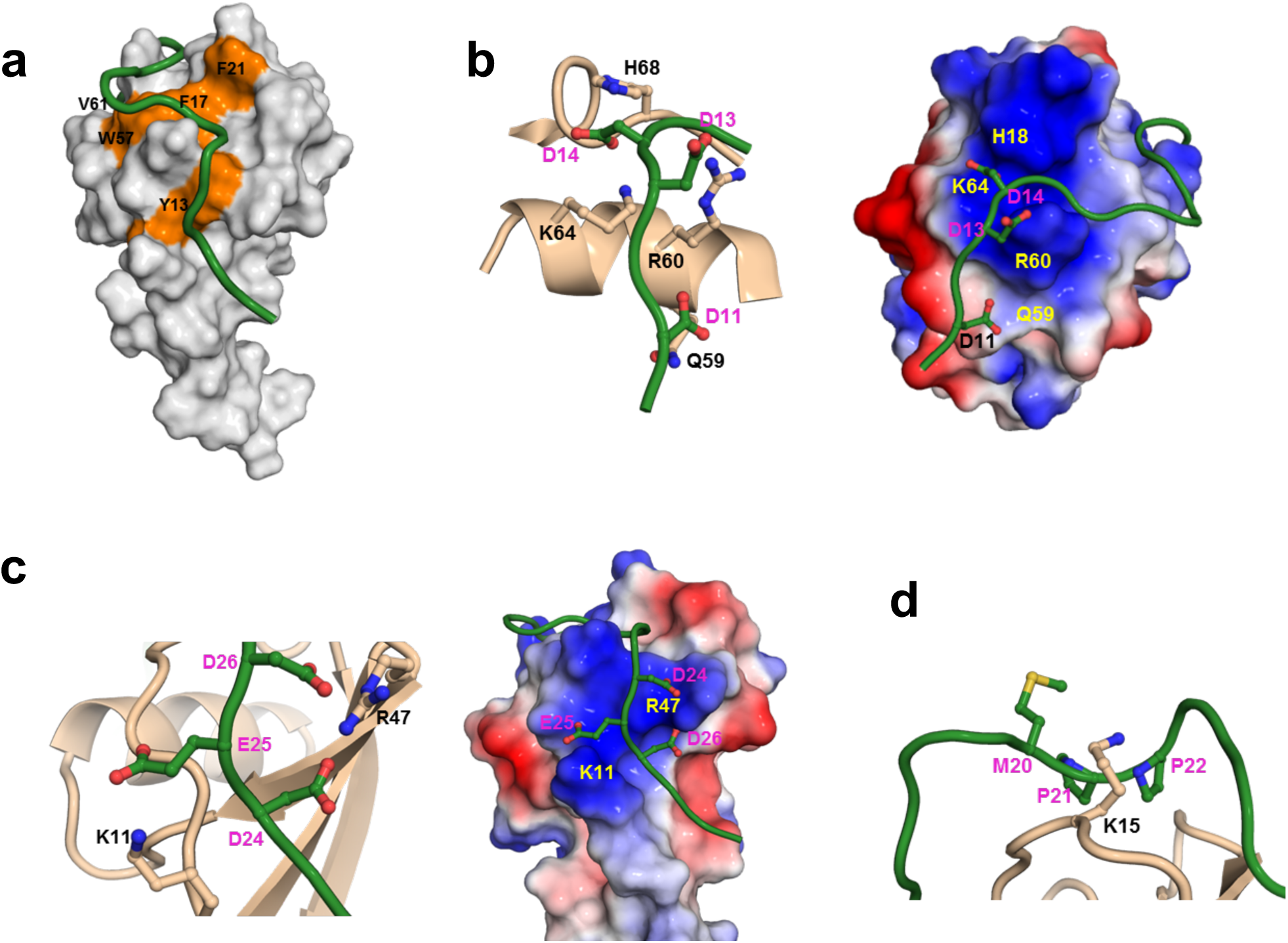
CXCL8-CXCR1 Site-I interactions. (**a**) Surface representation highlighting hydrophobic interactions in the CXCL8-CXCR1 N-domain complex. Key CXCL8 non-polar residues are labeled and shown in orange and CXCR1 is shown in green. (**b**) A schematic highlighting interactions of the first ionic cluster. CXCL8 residues are labeled in black and CXCR1 residues in magenta. Electrostatic representation of the first ionic cluster. CXCL8 residues are labeled in yellow and CXCR1 residues in magenta. (**c**) A schematic highlighting interactions of the second ionic cluster. CXCL8 residues are labeled in black and CXCR1 residues in magenta. Electrostatic representation of the second ionic cluster. CXCL8 residues are labeled in yellow and CXCR1 residues in magenta. (**d**) A schematic highlighting CXCL8 Lys15 side chain methylene groups packed against CXCR1 non-polar residues (shown as sticks and labeled in magenta).

### MD simulations of the CXCL8 monomer-CXCR1 complex

We set out to characterize how Site-I interactions determine binding at Site-II using MD simulation started from CXCL8 bound at Site-I. To this end, we first generated a structural model of CXCL8 monomer bound at Site-I of the CXCR1 receptor by merging the NMR structure of CXCL8-CXCR1 N-domain complex and a previously determined CXCR1 structure (Park et al., 2012). Using inter and intra molecular NOEs as distance constraints, we generated a structural model using HADDOCK (Supplementary Fig. 2). We then carried out MD simulation of the CXCL8-CXCR1 complex for 800ns. During the initial phase of the simulation, we used H-bonds derived from the NMR structure as distance restraints. The restraints were removed once stabilizing interactions with Site-II were formed.

The time evolution of the backbone r.m.s.d. demonstrates that the simulation has equilibrated after about 100ns (Supplementary Fig. 3). The largest structural reorganization occurred in the initial 50ns of the simulations and primarily involves the CXCR1 loops, while the structured regions of CXCL8 and the helices of CXCR1 are stable during the entire simulation. This can also be seen from the plots of secondary structure contents (Supplementary Fig. 4) and backbone root mean square fluctuations (r.m.s.f.; Supplementary Fig. 5). In contrast to the minimal fluctuation of the secondary structure elements, the termini of CXCL8 and the termini as well as the intra- and extracellular loops of CXCR1 experience large conformational fluctuations. As many of these flexible regions are involved in intermolecular interactions, the MD data provided several novel insights into how CXCL8 bound at Site-I searches and binds Site-II residues. In addition to ELR residues, several CXCL8 N-loop residues also bound at Site-II, indicating binding at two spatially distinct receptor sites are not discrete events but involves extensive crosstalk between the sites (Figs. 3-5; Supplementary Fig. 6). In particular, reorganization of Site-I interactions allow CXCL8 to swivel about the second ionic cluster, which increases the search radius for binding Site-II residues. We group CXCL8 residues on the basis of their interactions into four classes.

**Figure 3.**
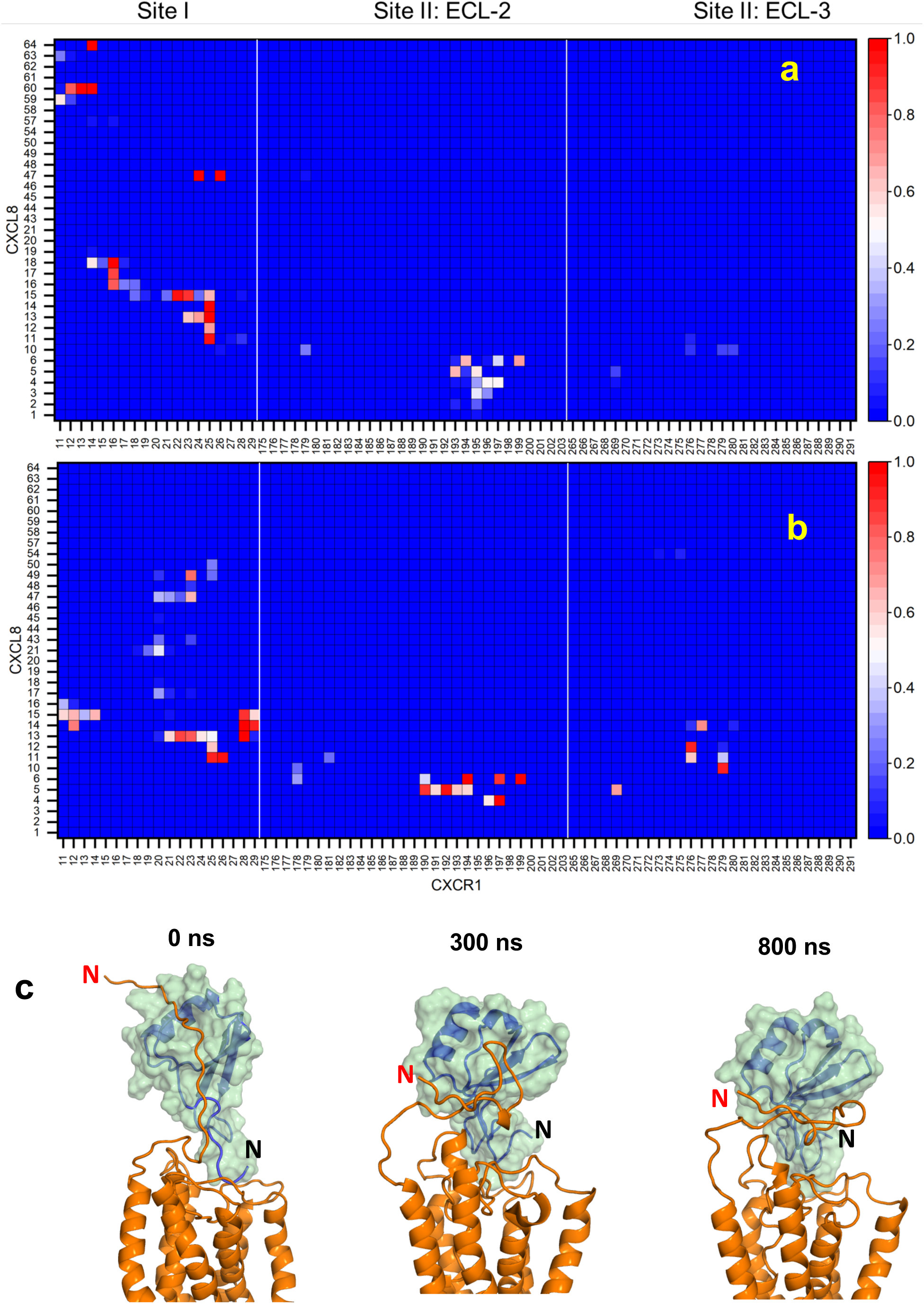
Molecular dynamics of the CXCL8-CXCR1 complex. Heat maps of the average contact frequency between all CXCL8 and CXCR1 residues during the first 50ns (**a**) and last 100ns (**b**). Only residues that come in contact during the simulations are shown. (**c**) Snapshots of the CXCL8-CXCR1 complex at different time points (0, 300, and 800ns) during the MD simulation. CXCR1 is shown in orange and CXCL8 is shown as both surface and ribbon representations in green. The N-terminus of CXCL8 and CXCR1 are labeled in red and black, respectively.

Group-I consists of residues Glu4, Leu5, Arg6, Ile10, His33, and Glu38. These residues (except Ile10), which are not involved in Site-I interactions, are now exclusively engaged in binding at Site-II. In the Site-I structure, ELR residues are unstructured and His33 is observed in the proximity of the CXCL8 N-terminal residues, and its imidazole side chain is in an environment that is different from the bulk solvent as inferred from pKa measurements (Sepuru and Rajarathnam, 2018). During the first 50 ns of the MD simulation, Glu4 and Arg6 are engaged in specific ionic interactions with ECL2 K197 and D194, respectively (Figs. 3, 4). These ion pairs are stable and retained throughout the MD run. Both D194 and K197 are unique to CXCR1 with the corresponding residues in CXCR2 being asparagines, indicating that Site-II binding interactions are different between the receptors. During the initial 50 ns, Leu5 is observed in the proximity of ECL2 N193 and T195, and subsequently binds to a pocket defined by ECL2 V190 and L191, and ECL3 R269 (Figs. 3, 4). Interestingly, His33 starts interacting with ECL3 R269 about the same time as Leu5, suggesting concerted interactions that are stable until the end of the MD run. The first three N-terminal residues (Ser1-Ala2-Lys3), which are not critical for activity (Clark-Lewis et al., 1993), are pointed away from the binding groove. In both the free and Site-I structures, the carboxylate group of Glu38 is H-bonded to Gln8 backbone amide proton (Fig. 3, Supplementary Fig. 7). During the MD simulations, Glu38 disengages in the first 10ns, and is observed to interact with R279 and R280 in the final MD structures. Ile10, which is packed against CXCR1 Y28 in the Site-I structure, disengages in the first 50 ns, and is initially observed in a pocket defined by ECL2 I279, ECL3 R279 and R280, and by the end of the run, is essentially packed against ECL3 R279 (Fig. 3).

**Figure 4.**
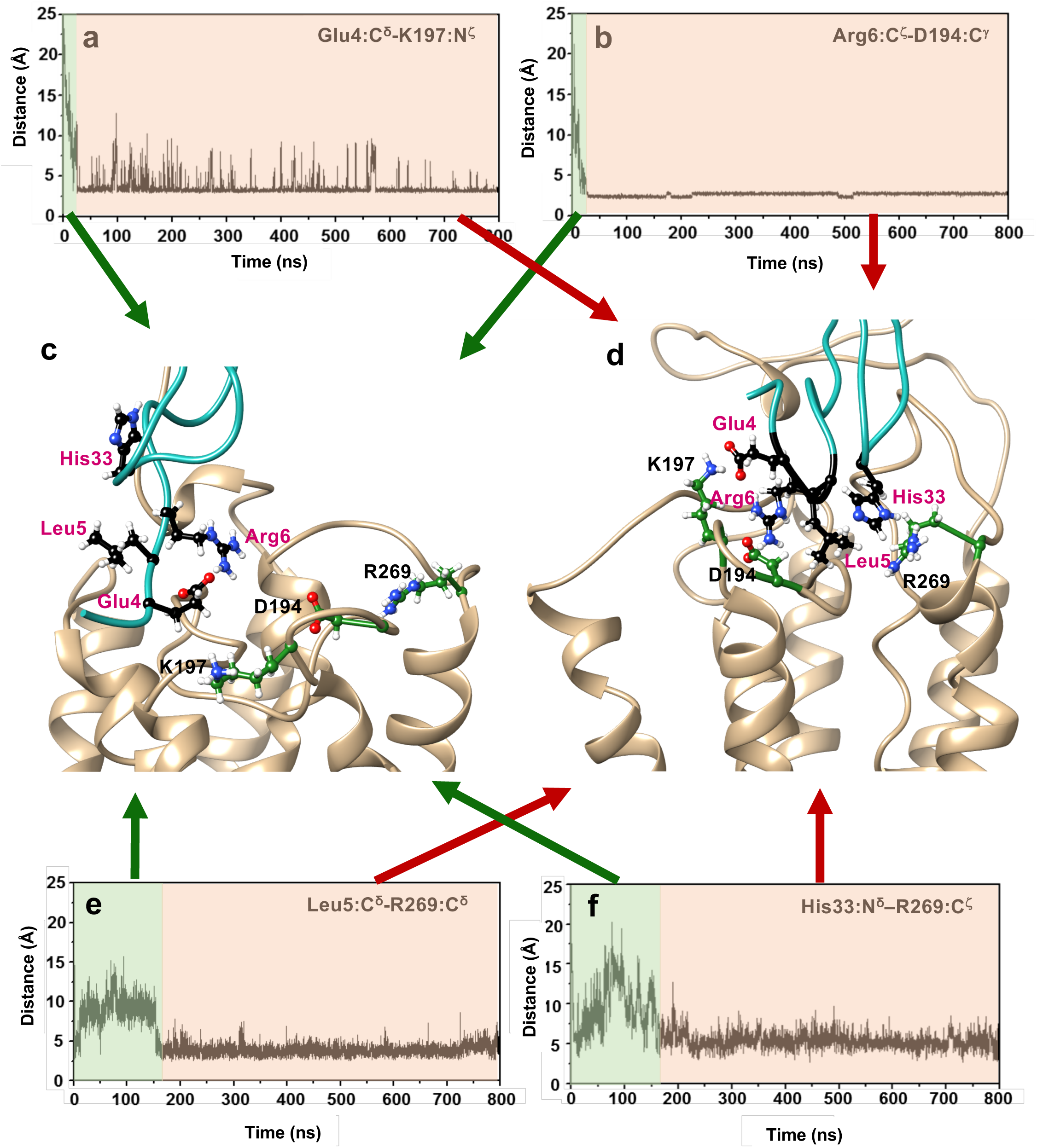
Site-II interactions. Distance plots between Glu4:C^δ^-K197:N^ζ^ (**a**), Arg6:C^ζ^-D194:C^γ^ (**b**), Leu5:C^δ^-R269:C^δ^ (**e**), and His33:N^δ^–R269:C^ζ^ (**f**). The plots shows that CXCL8 N-terminal ‘ELR’ and His33 residues recognize and bind their specific CXCR1 partners within 150ns. Structural snapshots of the CXCL8–CXCR1 complex at the beginning (**c**) and end (**d**) of MD snapshots, showing CXCL8 N-terminal ‘ELR’ and His33 residues (in red) recognizing their specific CXCR1 partners (in black).

Group-II consists of residues Lys11, Thr12, and Ser14. These residues, involved in ionic interactions in the NMR Site-I structure, undergo a reorganization and also start interacting with Site-II residues during the MD run. In the Site-I structure, Lys11 interacts with E25 and Y27, Thr12 interacts with E25 and D26, and Ser14 interacts with E25. Towards the end of the MD run, Lys11 interacts with E25 and D26 at Site-I and with S276 and R279 at Site-II; Thr12 retains its interaction with E25 at Site-I and now also interacts with S276 at Site-II (Supplementary Fig. 8); and Ser14 interacts with S28 and P29 at Site-I and with C277 at Site-II (Figs. 3, 5). Humans carrying a S276T mutation show impaired neutrophil function and are more susceptible to infection (Swamydas et al., 2016), highlighting the importance of these interactions for receptor function. These observations are also unprecedented, and make a compelling case for structural plasticity and dynamics of the binding residues in coupling Site-I and Site-II interactions.

**Figure 5.**
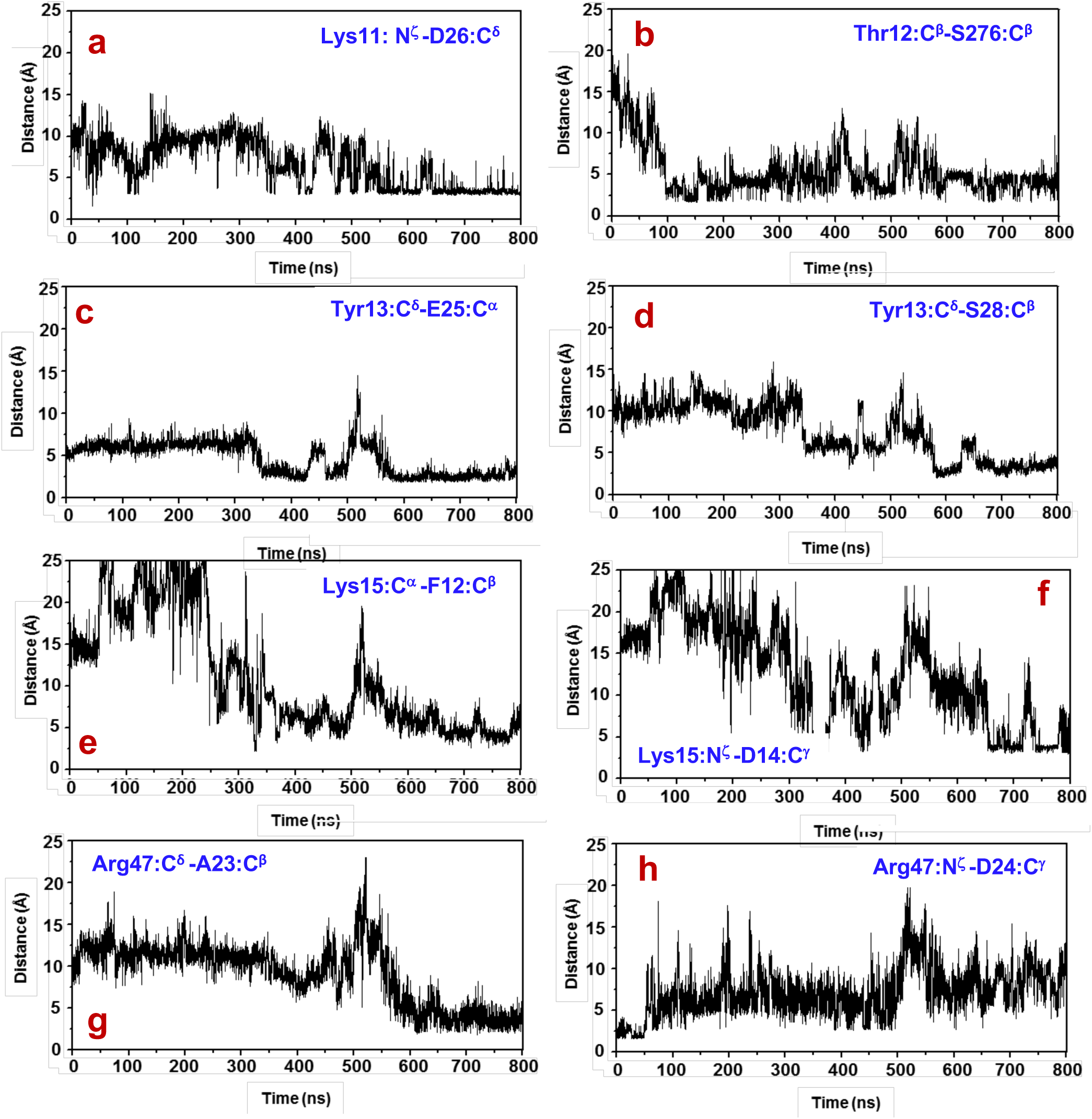
MD profiles of Site-II interactions. Distance plots of (**a**) Lys11:N^ζ^ -D26:C^γ^, (**b**) Thr12:C^β^-S276:C^β^, (**c**) Tyr13:C^δ^-E25:C^α^, (**d**) Tyr13:C^δ^-S28:C^β^, (**e**) Lys15:C^α^-F12:C^β^, (**f**) Lys15:N ^ζ^-D14:C^γ^, (**g**) Arg47:C^δ^-A23:C^β^, and (**h**) Arg47:N^ζ^–D24:C^γ^.

Group-III consists of residues Tyr13, Lys15, Phe17, Phe21, and Arg47. These residues are involved in ionic/packing interactions in the NMR Site-I structure, and undergo a reorganization during the MD run and are now engaged with different Site-I residues. In the Site-I structure, Tyr13 is packed against A23, D24, and E25; Lys15 is packed against M20, P21, and P22; and Arg47 is involved in ionic interactions with D24 and D26 (Fig. 2). During the MD run, Lys15 is found to be dynamic and its side chain spanning most of the binding interface from F12 to P29, and towards the end, is involved in ionic interactions with D13 and D14 (Figs. 3, 5). Tyr13 is also dynamic and interacts with P21, P22, A23, D24, and E25. During the MD run, Arg47 disengages from D24 and D26, and towards the end, is packed against M20, P21, P22, and A23. It is notable that interactions of Lys15 and Arg47 are switched between the NMR Site-I and final MD structures (Figs. 3, 5). Both Phe17 and Phe21 interact with N16 and P21 in the Site-I structures. Phe17 was observed to interact with several non-polar residues with longer residence times around M20 and P22, and Phe21 was observed to be packed against M20, P21, and P22 throughout the MD run (Fig. 3).

Group-IV consists of residues His18, Arg60, and Lys64. These residues are involved in ionic interactions with D11, D13, and D14 in the NMR Site-I complex, but disengage around 250ns and formed no new intermolecular interactions (Fig. 3). Therefore, Group-IV residues exclusively play a role in stabilizing Site-I interactions during the docking stage and play no role in Site-II interactions. Group-IV interactions result in CXCR1 N-terminal residues binding in an extended fashion as shown in Figs. 1 and 3c (at the beginning of the MD run). However, losing these interactions results in a more compact N-terminal domain (MD structures at 300 and 800 ns). MD structural snapshots reveal that Lys15 (Group-III) and its neighboring residues take the place of Group-IV residues for interacting with the acidic residues of the first ionic cluster (Fig. 3), CXCR1 A23 is packed against CXCL8 Tyr13, Arg47, and Leu 49, and that the stretch of CXCR1 residues from L15 to P22 undergo segmental motions with A23 and the first ionic cluster functioning as pivot points (Supplementary Fig. 9). These data also indicate that interactions of Lys15 are transient, suggesting the corresponding receptor residues are unlikely to be structured if a crystal structure of the CXCL8-CXCR1 were to become available.

## Discussion

Chemokine N-terminal ELR and N-loop residues, and 30s-loop, β_3_-strand, and C-helical residues that are in the proximity of ELR and N-loop residues identified in the current MD study mediate receptor function (Clark-Lewis et al., 1993, 1994; Kuschert et al., 1998; Lowman et al., 1996; Schraufstatter et al., 1995; Suetomi et al., 1999). We have previously shown that the conserved disulfides and the CXC motif that are in the proximity of ELR and N-loop residues are also critical for function (Joseph et al., 2010; Joseph et al., 2013b; Rajagopalan et al., 2007; Rajarathnam et al., 1999). N-terminal, N-loop, 30s-loop, disulfides, and the CXC motif are also conformationally dynamic, suggesting that cross-talk between these residues play an important role for function. We had proposed a ‘conformational-selection’ model, whereby the CXCL8 N-terminal, N-loop and 30s-loop, and the disulfides and as well as chemokine receptors exist as a conformational ensemble, such that their association occurs via a series of coordinated conditional and discrete binding steps (Joseph et al., 2013b). The structural plasticity and reorganization of Site-I interactions observed in the current MD run are in agreement with such a model.

Residue-specific dynamic measurements of the CXCL8-CXCR1 N-domain complex indicate that N-loop residues - Ile10, Thr12, Tyr13, Ser14, and Lys15 - undergo chemical exchange at slower millisecond-microsecond (ms-μs) time scales (Joseph et al., 2018). Chemical exchange for a set of contiguous N-loop residues provides a structural basis for how a coupled network that is dynamically primed for conformational selection could mediate Site-II interactions. The observation that Ile10 belongs to Group-I, Thr12 and Ser14 belong to Group-II, and Tyr13 and Lys15 belong to Group-III is striking, providing crucial insights into how structural plasticity and conformational dynamics of these residues are integral to the search process and binding Site-II residues. Ile10Ala mutant is substantially impaired for receptor activity but shows only two-fold reduced affinity for the CXCR1 N-domain (Clark-Lewis et al., 1994; Ravindran, 2010), suggesting that the steric bulk of Ile10 is less important for the initial docking but more important for binding at Site-II. The residue corresponding to Ile10 tends to be non-polar or aromatic in both CXC and CC chemokines (Supplementary Fig. 10), suggesting that this residue in other chemokines plays a similar role in coupling Site-I and Site-II interactions.

Our MD data indicate that binding Site-II is not a simple event but is coupled to a complex reorganization of Site-I interactions (Figs. 3 to 5). The observation that these changes occur relatively fast (mostly within 250ns) also corroborates a pivotal role for conformational plasticity and that crosstalk between Site-I and Site-II is a highly dynamic process. A previous MD study using CXCL8 bound to CXCR1 supplemented with a homology modeled CXCR1 N-domain based on the bovine rhodopsin structure could not describe interactions of the ELR residues (Liou et al., 2014). This highlights the importance of a Site-I structure that captures native interactions for characterizing how binding at Site-I determines Site-II interactions.

CryoEM structure of CXCL8 bound to CXCR2 in the presence of G protein (active form) was recently reported (Liu et al., 2020). The structure reveals that Glu4 is engaged in ionic interactions with three arginines located in ECL2 and ECL3, Leu5 is involved in packing interactions with two valines in the ECL2, and Arg6 forms a H-bond with the carbonyl oxygen of a threonine in ECL3. These interactions are different from those observed in our MD study of the CXCL8-CXCR1 complex. Glu4 and Arg6 are engaged in specific ionic interactions with a ECL2 lysine and aspartate (Figs. 3, 4). A conservative mutation of Arg6 to a lysine abolishes receptor activity (Clark-Lewis et al., 1993). Therefore, Arg6 interacting with a backbone carbonyl in the CXCL8-CXCR2 structure is unexpected considering the absolute requirement of the arginine guanidinium side chain for affinity and activity. The observed differences could be due to differences in CXCR1 vs. CXCR2 Site-I interaction or that CXCL8 binds to a different subset of Site-II receptor residues in the active vs. inactive forms. Note that our MD study was of CXCL8 binding to CXCR1 in the inactive form whereas the cryoEM structure was of CXCL8 binding to CXCR2 in the active form.

Comparing the CXCL8-CXCR1 N-domain structure to CXCL12-CXCR4, CCL5-CCR2, and CCL11-CCR1 N-domain Site-I structures show some similarities but also notable differences (Abayev et al., 2018; Millard et al., 2014; Veldkamp et al., 2008; Ziarek et al., 2017) (Fig. 6). Structures reveal that chemokines bind the most proximal receptor N-terminal residues up to the preceding residue of the conserved cysteine (Fig. 6E). This is notable considering receptor N-domains show low sequence homology that is evident even for related receptors CXCR1 and CXCR2 (Supplementary Fig. 11). Receptor N-domain sequence lengths vary, are acidic due to many Asp/Glu, and in addition, receptor N-domain tyrosines are also known to be sulfated (Ludeman and Stone, 2014). In the CXCL8-CXCR1 N-domain Site-I structure (Fig. 1), proximal N-domain residues (D11 to P29) mediate binding, whereas the distal N-domain residues are unstructured. However, in the CXCL12-CXCR4 N-domain structure, besides the CXCR4 proximal N-terminal residues (N11-K25), distal N-terminal residues (S5-D10) are also involved in binding (Ziarek et al., 2017). This set of distal N-terminal residues bind to CXCL12 β_1_ residues, and has been labeled as Site 0.5 on the basis of a similar observation for CXCL12 binding to ACKR3 (Gustavsson et al., 2017). Site 0.5 interactions are not observed in our CXCL8-CXCR1 N-domain structure or in the other two CC chemokine Site-I structures, indicating these interactions come into play only for some chemokine-receptor pairs. The structures of CCL5-CCR5 and CCL11-CCR1 N-domain Site-I complexes show that fewer receptor residues mediate binding and also that these interactions involve different regions of the chemokine (Figs. 6C and 6D). Solution NMR studies have shown that the N-domain of the intact CXCR1 is in equilibrium with the membrane, and that only the free N-domain can bind the ligand (Park et al., 2011). Using time-resolved fluorescence, surface pressure, and red edge excitation shift measurements, it has also been shown that the CXCR1 N-domain interacts with the membrane surface (Haldar et al., 2010). These observations provide a glimpse of the diversity in Site-I interactions, and also suggesting that coupling between Site-I and Site-II interactions will accordingly vary.

**Figure 6.**
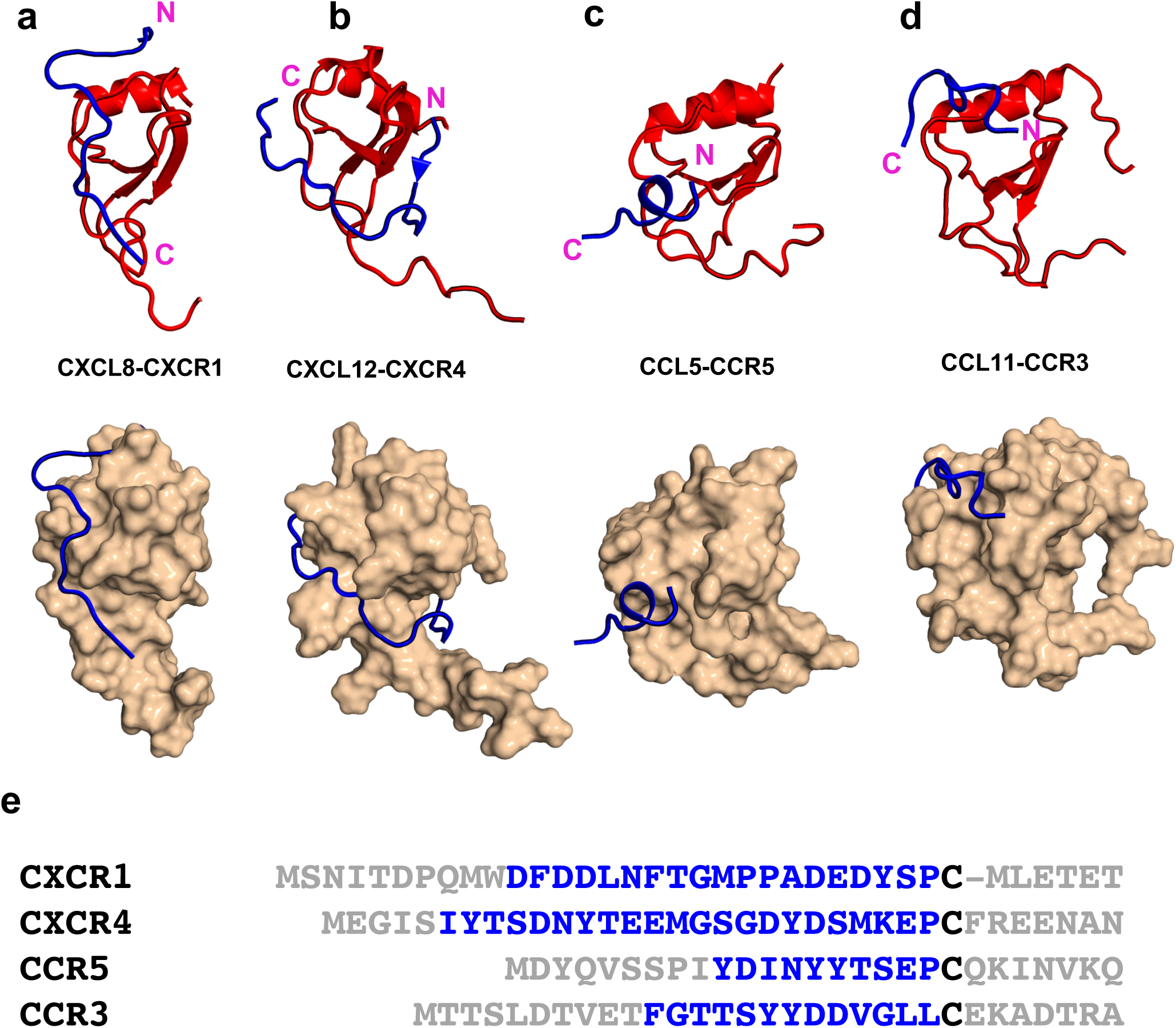
Comparison of chemokine-receptor N-domain site-I structures. (**a**) CXCL8-CXCR1 (PDB ID: 6XMN), (**b**) CXCL12-CXCR4 (PDB ID: 2N55), (**c**) CCL5-CCR5 (PDB ID: 6FGP), and (**d**) CCL11-CCR3 (PDB ID: 2MPM). The structures were aligned pairwise against CXCL8-CXCR1 structure. The top and bottom panels show ribbon and surface representations, respectively. The chemokine receptor N-domains are shown as a ribbon (blue). (**e**) Sequences of receptor N-domain. Residues involved in binding are in blue and the conserved cysteine is in black.

A solution structure of a CXCR1 N-domain peptidomimetic bound to dimeric CXCL8 is available (Skelton et al., 1999). The peptidomimetic corresponds to residues 9 to 29 containing a 6-amino hexanoic acid linker in place of residues 15-19. Structures reveal significant differences, as evident by NOEs from 20 CXCL8 and 17 CXCR1 residues for the monomer and from only 6 CXCL8 and 7 CXCR1 residues for the dimer (Supplementary Fig. 12). CXCR1 residues P21 to P29 bind in an extended fashion in the monomer-bound structure, whereas these residues adopt a more compact structure in the dimer-bound form (Supplementary Fig. 12). Backbone relaxation and amide exchange measurements show that the monomer is more dynamic (Grasberger et al., 1993; Joseph et al., 2013a; Joseph et al., 2018; Rajarathnam et al., 1995; Rajarathnam et al., 1994), indicating conformational fluctuations are critical for the higher affinity and extensive binding interactions observed for the monomer. Fewer interactions for the dimer compared to the monomer have also been observed in CXCL12 binding to the CXCR4 N-domain (Veldkamp et al., 2008; Ziarek et al., 2017).

Our MD data for the CXCL8-CXCR1 complex indicate that several Site-I interactions are transient, residues that mediate Site-I interactions continue to be dynamic even after Site-II is engaged, and structural plasticity allows the same residue binding at both sites. CryoEM and crystal structures of CXCR2 bound to CXCL8, CCR6 bound to CCL20, CCR5 bound to CCL5 antagonist (5P7-CCL5), CXCR4 bound to vMIPII (a promiscuous viral chemokine antagonist), and a constitutively active viral chemokine receptor US28 bound to CX_3_CL1 are known (Burg et al., 2015; Liu et al., 2020; Qin et al., 2015; Wasilko et al., 2020; Zheng et al., 2017) (Fig. 7). The coordinates of the CXCL8-CXCR2 complex are yet to be released and the Site-I structure of the CXCL8-CXCR2 N-domain is not known. Comparison of NMR CCL5-CCR5 N-domain complex and 5P7CCL5-CCR5 crystal structures reveals that several N-domain residues that are structured in the former are unstructured in the latter (Fig. 7F). Structures of CCL20-CCR6, vMIPII-CXCR4, and CX3CL1-US28 complexes reveal electron density for only a few N-domain residues. Site-I structure of vMIPII-CXCR4 N-domain is not known but the structure of the CXCL12-CXCR4 N-domain shows extensive Site-I interactions. These observations collectively indicate that conformational dynamics of receptor N-domain residues preclude these residues from being ‘visible’ in the crystal and cryoEM structures, and also suggest that loss and reorganization of Site-I interactions on binding Site-II could be a basic mechanism underlying ligand recognition by all chemokine receptors.

**Figure 7.**
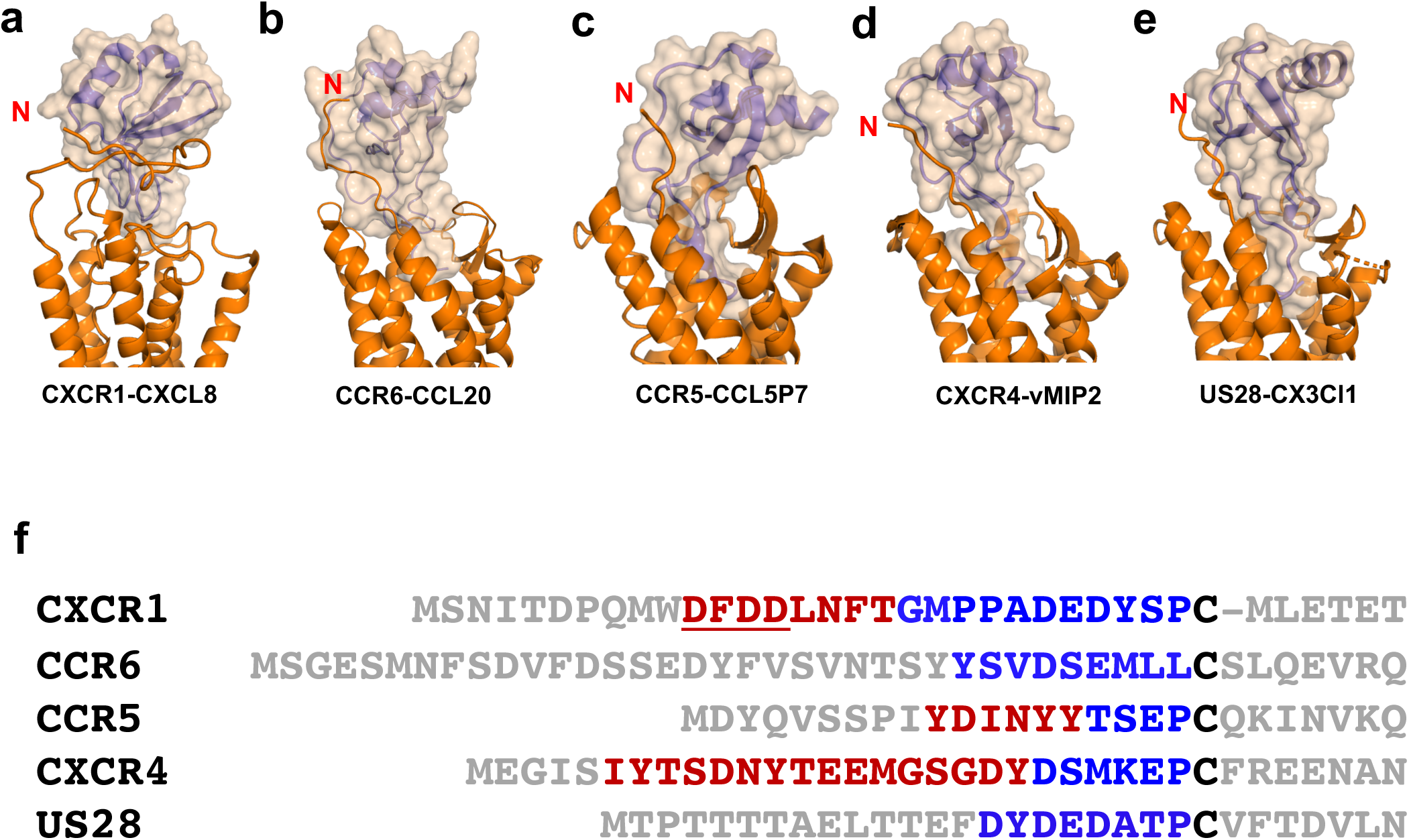
Comparison of chemokine-receptor complex structures. **(a)** CXCL8-CXCR1; (**b**) CCL20-CCR6 (PDB ID: 6WWZ); (**c**) CCL5P7-CCR5 (PDB ID: 5UIW); (**d**) CXCR4-vMIP2 structure (PDB ID: 4RWS); and (**e**) US28-CX3Cl1 (PDB ID: 5WB2). The receptors and chemokines are shown in orange and purple, respectively. The receptor N-termini are denoted in red. **(f)** Sequences of receptor N-terminal domain. Residues involved in binding are in blue, and residues involved in binding in the NMR Site-I structures but not visible in the crystal structures are in red. The ‘DFDD’ residues of CXCR1 are underlined as MD studies indicate that these residues are involved in transient interactions and so unlikely to be observed in a crystal structure.

Our MD data show that the first three CXCL8 N-terminal residues are not located in the groove, and residues Glu4-Leu5-Arg6 interact with EC loop residues. ELR-CXC chemokines have as few as one (CXCL7) and as many as nine (CXCL5) residues preceding the ‘ELR’ motif (Supplementary Fig. 11), suggesting these residues finetune Site-II interactions. The cryoEM structures of CXCL8-CXCR2 and CCL20-CCR6 also reveal that the CXCL8 and CCL20 N-terminal are located closer to the surface (Liu et al., 2020; Wasilko et al., 2020). However, crystal structures of 5P7CCL5-CCR5 and vMIPII-CXCR4 complexes reveal that the chemokine N-terminal residues are located deep in the TM groove and are in the proximity of toggle switch residues that trigger the conformational change for receptor activation (Wescott et al., 2016). These observations suggest that there is no absolute requirement that chemokine residues must be located in the proximity of toggle-switch residues for receptor activation.

The past decade has seen an explosion of GPCR including chemokine receptor structures that can be attributed to advances in receptor expression and folding, reconstitution with detergents, and constructs that are stable and more amenable to crystallization (Kruse et al., 2013; Rosenbaum et al., 2007). The latter technology includes replacing a flexible IC loop with T4 lysozyme, using nanobodies to stabilize a specific conformer, and/or solving the structure bound to agonists, antagonists, and/or small molecule inhibitors that were critical for determining chemokine receptor structures and generating structural models (Bhusal et al., 2020; Wu et al., 2010; Zheng et al., 2016). These structures represent snapshots and are not sufficient for providing a spatiotemporal description of how the chemokine bound at Site-I searches and binds the residues at Site-II. In this study, we show that a hybrid NMR-MD approach can provide this information and that long-range coupled motions between Site-I and Site-II orchestrate CXCL8 monomer binding to the CXCR1 receptor. On the basis of our current studies and available literature, we propose tunable coupled motions between Site-I and Site-II underlie the range of affinity, potency, and specificity observed for various chemokine-receptor pairs. We conclude that NMR-MD approach in complement with ever increasing number of structures, can provide molecular details that are crucial for understanding the basic principles that govern chemokine-receptor interactions and their roles in human pathophysiology, thus paving the way for designing highly specific and potent drugs.

## Materials and Methods

### Protein Expression and Purification

CXCL8(1–66) monomer and CXCR1 N-domain (1-29) peptide were cloned, expressed, and purified as described previously (Fernando et al., 2007).

### Chemical shift assignments and structure calculation

Chemical shift assignments of CXCL8 and CXCR1 N-domain in the bound form were obtained using CBCA(CO)NH, HNCACB, CC(CO)NH, ^15^N-edited TOCSY-HSQC, and HCCH-TOCSY experiments using 0.4 mM ^13^C/^15^N-CXCL8 bound to unlabeled CXCR1 N-domain and unlabeled CXCL8 bound to ^13^C/^15^N-CXCR1 N-domain samples (Gardner and Kay, 1998). The spectra were collected at 303K on a Bruker 800 MHz spectrometer with a cryogenically cooled probe head. The samples were prepared in 50 mM phosphate pH 6.0 containing 10% D_2_O and 0.01% sodium azide. The backbone dihedral angles were predicted using the TALOS software (Shen et al., 2009). The chemical shifts were referenced using DSS as the internal standard. The NMR data were processed using Bruker Topspin and analyzed by Sparky. Intramolecular distance restraints were obtained from ^15^N-edited NOESY-HSQC and ^13^C-edited NOESY-HSQC experiments optimized either for the aliphatic or the aromatic region. The distance constraints and angular restraints for CXCL8 alone were used to calculate structures of CXCL8 and CXCR1 N-domain in the complex using ARIA/CNS suite of programs (Rieping et al., 2007). Intermolecular NOE distance restraints were obtained from ^13^C (ω_2_)-edited/^12^C (ω_3_)-filtered NOESY-HSQC spectra of ^15^N/^13^C-CXCL8 monomer complexed to 3-fold excess of unlabeled CXCR1 N-domain and *vice versa*. Structure of the CXCL8-CXCR1 N-domain complex was calculated iteratively using distance, dihedral angle, and H-bonding restraints as input using ARIA/CNS and the PARALLHDG 5.3 force field. The quality of the calculated structures was assessed using PROCHECK.

### Model of CXCL8 bound at CXCR1 N-domain

A model of CXCL8 bound at N-domain of the full length CXCR1 receptor was generated using HADDOCK as follows. Initially, the missing N-terminal residues 1 to 28 were appended to the CXCR1 structure (PDB ID: 2LNL; residues 29-324) using PyMOL. For Site-I interactions, all experimental intramolecular and intermolecular NOEs were used as unambiguous restraints. For Site-II interactions, constraints between CXCL8 N-terminal Glu4 and Arg6 and all CXCR1 ECL2 and ECL3 acidic and basic residues were used as ambiguous restraints. A total of 4000 structures were generated using rigid body docking in HADDOCK (Dominguez et al., 2003). The top 100 structures that had the best intermolecular energies were then subjected sequentially to semiflexible simulated annealing and explicit solvent refinement during which the CXCL8 N-terminal ‘ELR’ and CXCR1 ECL2 and ECL3 acidic and basic residues were allowed to move freely. The pair-wise “ligand interface root-mean-square deviation matrix” over all structures was calculated, and final structures were clustered using a cutoff value of 6.0 Å. The clusters were sorted using r.m.s.d. and “HADDOCK score” (the weighted sum of a combination of energy terms), and the best scored structure was used as the initial model for MD studies.

### Molecular dynamics simulations

We embedded the modeled CXCL8 monomer-CXCR1 complex structure into a fully hydrated bilayer consisting of 120 16:0/18:1 phosphatidylcholine (POPC) lipids per leaflet, using CHARMM GUI (Jo et al., 2008). The size of the water box was x = 93.6Å, y = 93.6Å, z = 113.0Å such that the minimum distance of a protein atom from the edge of the box was 10Å. The protein was solvated with TIP3P waters. The system was charge-neutralized and the ionic strength was set to 150mM by adding KCl. The final size of the system was 101,509 atoms. The protein-bilayer-water system was energy minimized for 2000 steps with lipid and protein heavy atoms fixed and equilibrated first for 200ps and then for four 100ps equilibration steps with a gradually decreasing restraint force constant. The equilibrated system was simulated with a 2fs time-step using SHAKE (Ryckaert, 1977) to restrain all bonds involving hydrogens. The simulation was conducted using the CHARMM36m force field (Huang et al., 2017) and the latest version of the NAMD program (Phillips et al., 2005).

We analyzed the simulations by calculating r.m.s.d. for the backbone atoms of the complex and the TM helices of CXCR1 after aligning using the C_α_ atoms of the TM helices, and those of CXCR1 and CXCL8 were calculated following alignment on the respective initial structures. To evaluate the conformational fluctuations of the complex, we calculated time-averaged r.m.s.f. of individual CXCR1 and CXCL8 residues after backbone alignment to the average structure (Zhou et al., 2017). We identified the interface residues by calculating the contacts between the heavy atoms of CXCR1 and CXCL8, with two residues were defined to be in contact if any of their atoms are within 4Å of each other. Using the contact information, we plotted a contact-map of the CXCR1-CXCL8 interface and calculated the time-averaged contact frequencies for individual residues.

## Author contributions

K. R., A. G., K.M.S., and V.N. designed research.

K.M.S. performed NMR experiments and structure determination under supervision of K.R.

V. N. and P.P. performed the MD simulations and analyses under supervision of A. G.

K. R. drafted the manuscript with contributions from A.G. and K.M.S. All authors reviewed the final manuscript.

## Acknowledgments

We thank Dr. Wang for NMR technical support. The authors acknowledge the Sealy Center for Structural Biology and Molecular Biophysics at the University of Texas Medical Branch at Galveston for providing research resources, and Texas Advanced Computing Center (TACC) and the Extreme Science and Engineering Discovery Environment (XSEDE grant No MCB150054) for providing computational resources. This work was supported in part by a grant from the National Institute of Health R01GM124233 to A.A.G. and a pilot grant from the Institute for Human Infections and Immunity at UTMB to K.R. V.N. is supported by UTHealth Innovation for Cancer Prevention Research Training Program Pre-Doctoral Fellowship (Cancer Prevention and Research Institute of Texas grant RP160015).

## Data availability

The solution NMR ensemble and related data sets of the CXCL8-CXCR1 N-terminal domain complex have been deposited in the Protein Data Bank (PDB) and Biological Magnetic Resonance Bank (BMRB) database under the codes PDB 6XMN and BMRB 50331, respectively.

