## supplementary figs. and tables for "Long-range coupled motions underlie ligand recognition by a chemokine receptor"

**Supplementary Figure 1.** **(a)** All atom and **(b)** backbone representations of the ensemble structures of the CXCL8-CXCR1 N-domain complex. CXCL8 and CXCR1 are shown in black and red, respectively.

**Supplementary Figure 2**. Ribbon representation of the CXCL8-CXCR1 complex generated using HADDOCK. Structures are shown at different orientations to highlight Site-I and Site-II interactions. CXCR1 is shown in purple and CXCL8 in teal.

**Supplementary Figure 3.** Time evolution of the backbone r.m.s.d. from the initial structure for all residues of the CXCR1-CXCL8 complex (black), CXCR1 (red) and CXCL8 (green), as well as for only the TM helices of CXCR1 (blue). The black dashed line separates the initial 50ns restrained portion of the simulation from the subsequent unrestrained 750ns.

**Supplementary Figure 4.** Time evolution of the secondary structure content in CXCR1 (top) and CXCL8 (bottom).

**Supplementary Figure 5.** Residue-based r.m.s.f. of the backbone atoms of CXCR1 (left) and CXCL8 (right) averaged over unrestrained (last 750ns) portion of the simulation, showing substantial fluctuations at the chain termini and the intracellular and extracellular loop regions.

**Supplementary Figure 6.** (**A-C**) An ensemble of the MD structural snapshots of the CXCR1-CXCL8 complex taken every 100ns. The ensemble is shown in different orientations to highlight the evolution of Site-I and Site-II interactions during the MD run. CXCR1 is shown in gray and CXCL8 in green. The ensemble was aligned on the backbone atoms of the TM helices of CXCR1. (**D-F**) An ensemble of structural snapshots at 0ns (red), 250ns (green), and 800ns (blue), that highlight coupling between Site-I and Site-II interactions. A single representative CXCR1 structure at 250ns is depicted in gray. (**G**) A heat map of the average contact frequency between all CXCL8 and CXCR1 residues during the entire 800ns MD simulation.

**Supplementary Figure 7. Intramolecular CXCL8 Gln8-Glu38 H-bonding interaction**. A schematic showing the fate of Gln8-Glu38 H-bond during the MD simulation. The initial structure at 0ns shows evidence for H-bond that is not present in the final 800ns structure. CXCL8 is shown in green and CXCR1 N-domain in purple. Glu38 carboxylate oxygens and Gln8 backbone amide are shown in red and blue, respectively.

**Supplementary Figure 8. CXCL8 Thr12-CXCR1 S276 interaction.** A distance plot showing H-bond formation between CXCL8 Thr12 and CXCR1 S276 (middle). The MD structures shows that these residues are far apart at initial 0ns (top) and are in H-bonding distance in the final 800ns structure (bottom).

**Supplementary Figure 9. Structural plasticity of Site-I interactions.** (**a**) An ensemble of the structural snapshots from 675ns to 800ns highlighting CXCL8 Lys15 (blue) interacting with CXCR1 N-domain. CXCR1 residues D11 to D14 are shown in orange, L15 to P22 in rainbow colors, and A23 in mauve. The ensemble is shown in different orientations to highlight concerted motion of L15 to P22 about A23 and D11 as pivot points. The individual time points (panels **b** to **g**) highlight stable interactions between CXCR1 A23 and CXCL8 Tyr13, Arg47 and Leu49, and transient interactions between CXCL8 Lys15 and CXCR1 D11 and D14.

**Supplementary Figure 10. Sequences of human CXC and CC chemokines.** Conserved cysteines and the residue corresponding to Ile10 are highlighted in green and red, respectively. The N-terminal 'ELR' residues are highlighted in blue.

**Supplementary Figure 11. Sequences of human CXC and CC receptor N-domains.** Conserved cysteine is highlighted in green, and aspartates (D) and glutamates (E) are highlighted in red.

**Supplementary Figure 12**. Comparison of CXCR1 N-domain bound to CXCL8 monomer (**a**) and CXCL8 dimer (**b**) structures. An ensemble of CXCR1 N-domain residues P21 to P29 are shown with side chains as sticks. A schematic of the intermolecular NOE contacts in the monomer-bound (**c**) and dimer-bound (**d**) complexes. Basic and acidic residues are highlighted in blue and red, respectively.

**Supplementary Table-I.** Structural Statistics for the NMR CXCL8-CXCR1 N-domain complex structures.

**Supplementary Table-II.** List of intermolecular NOEs that were used in the structure calculation.

**Supplementary Figure 1**

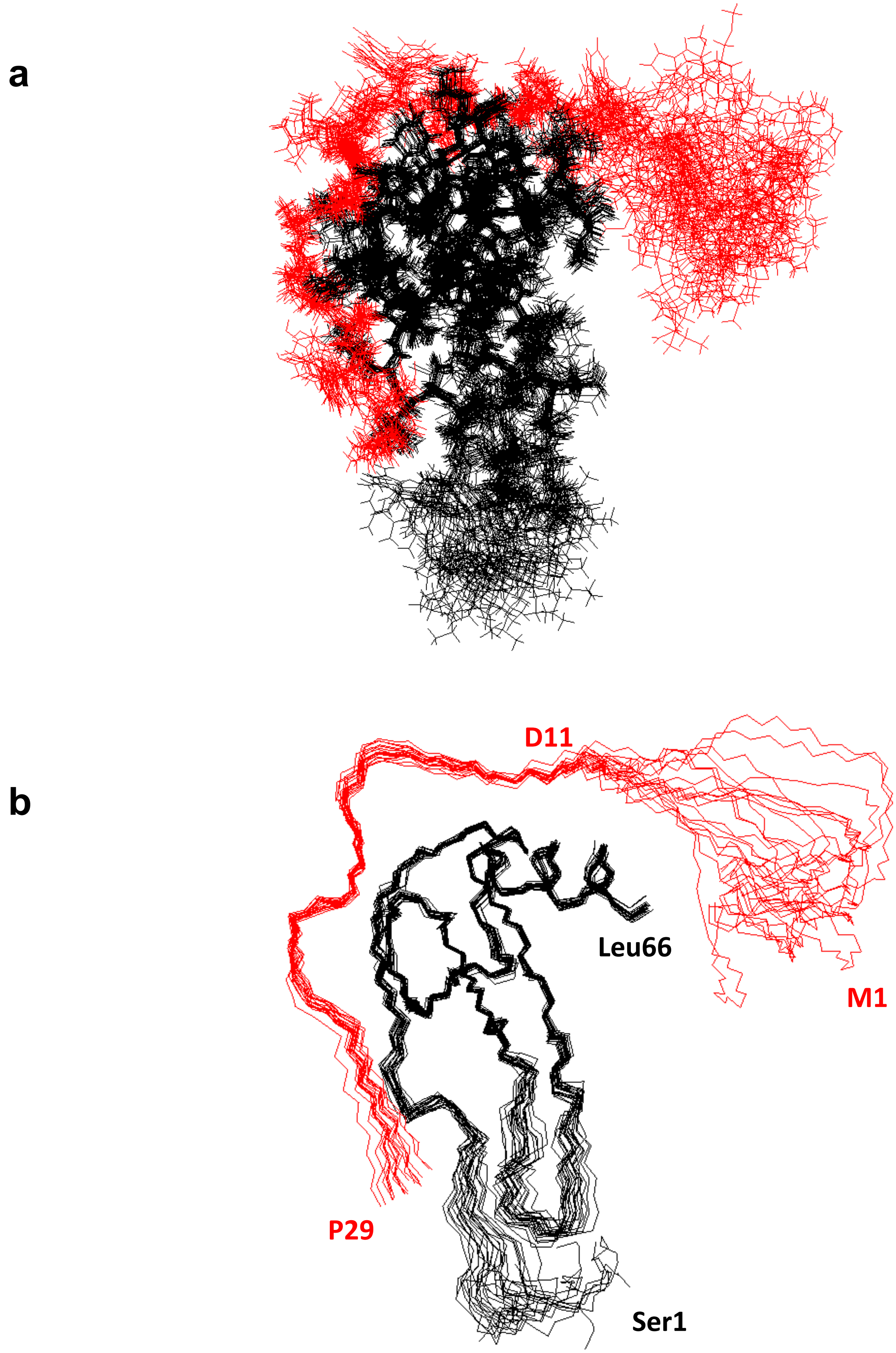

### Supplementary Figure 2

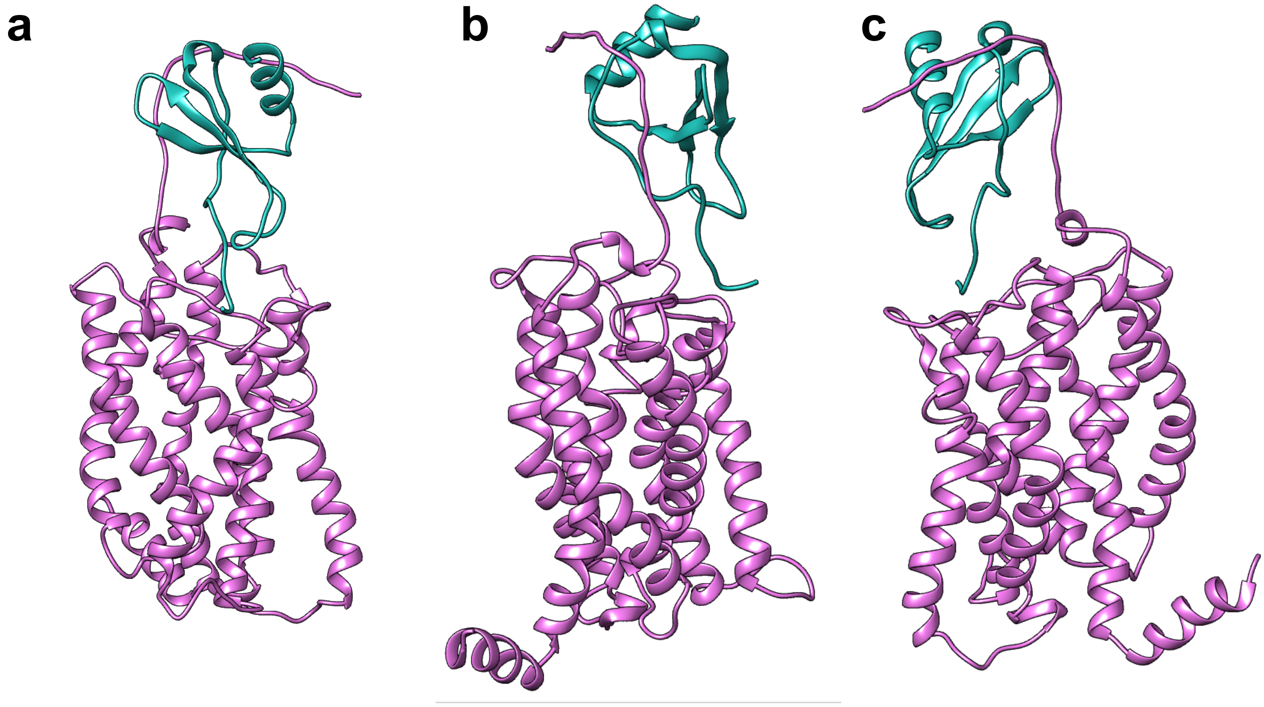

**Supplementary Figure 3**

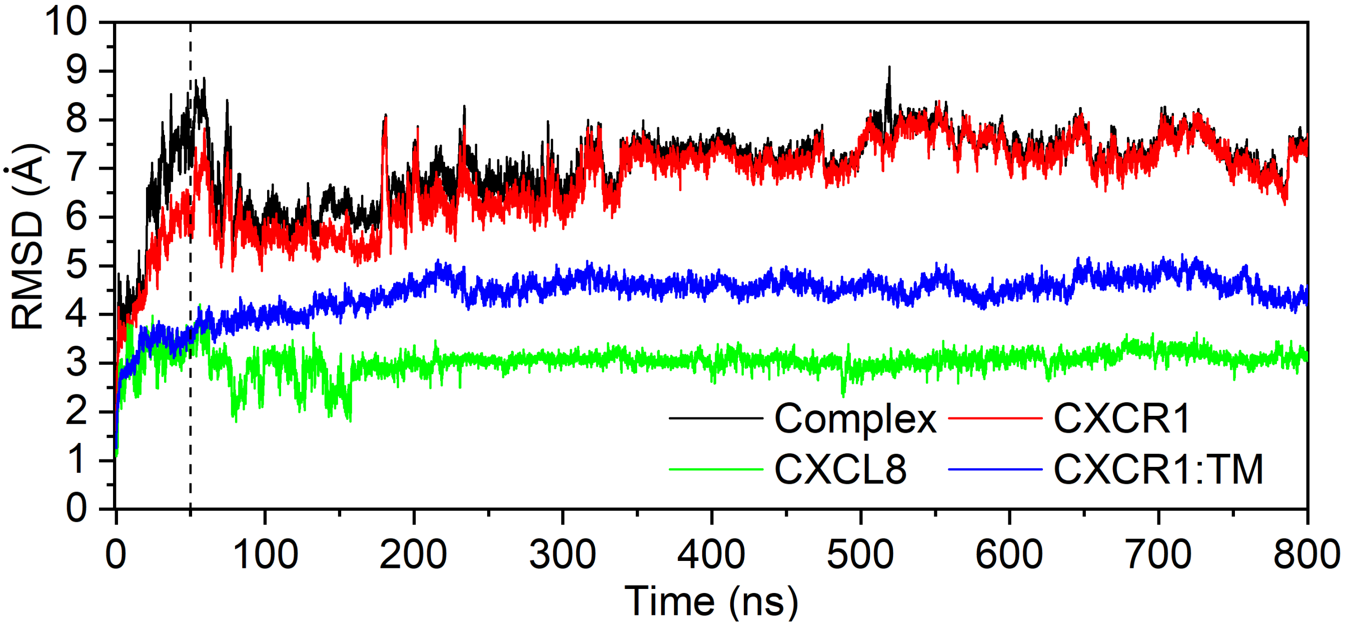

#### Supplementary Figure 4

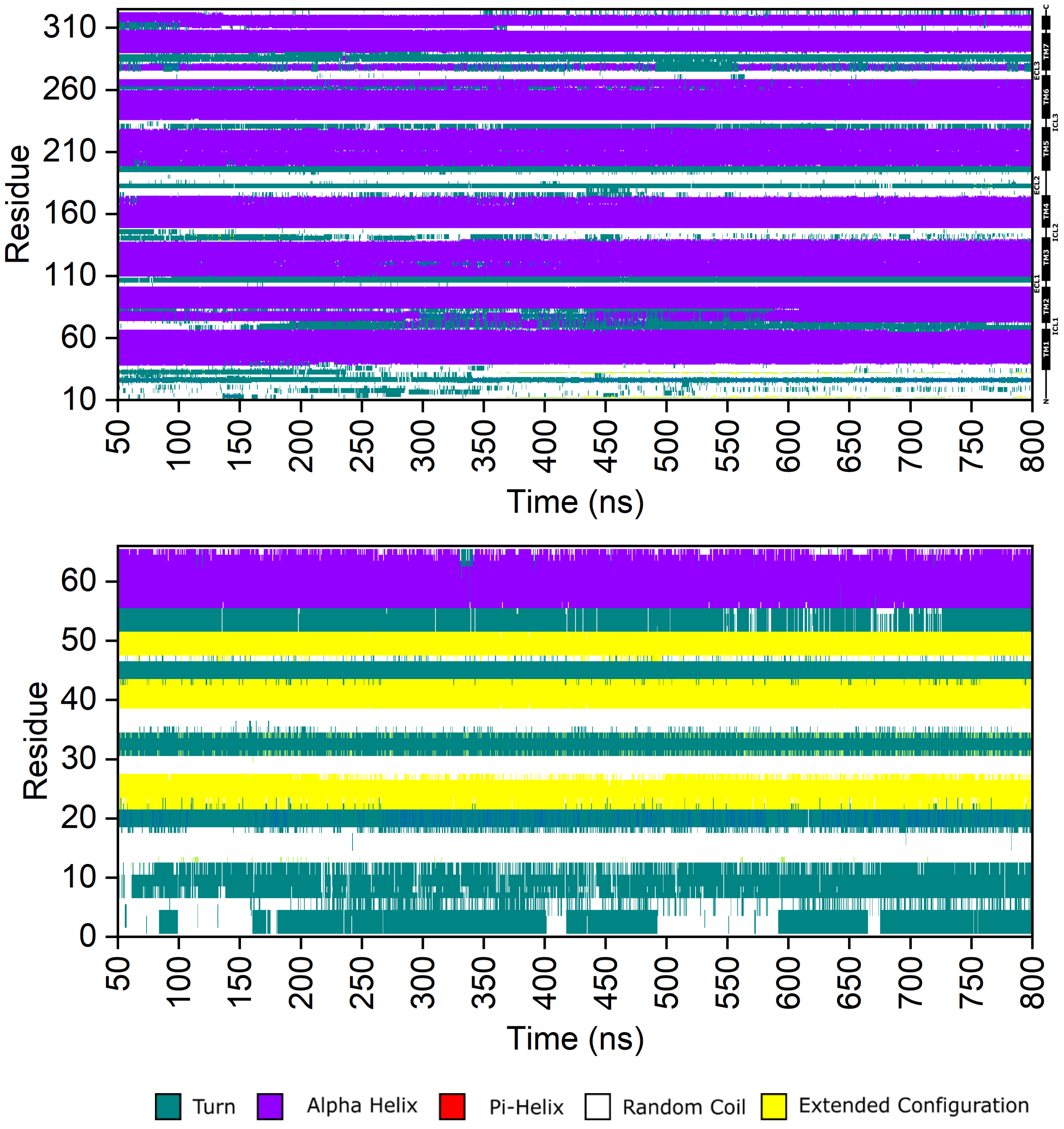

### Supplementary Figure 5

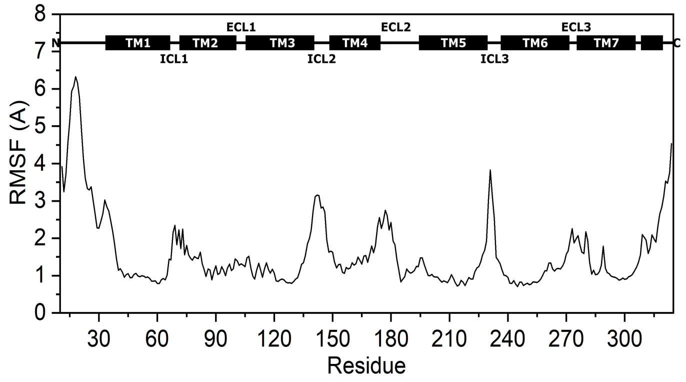

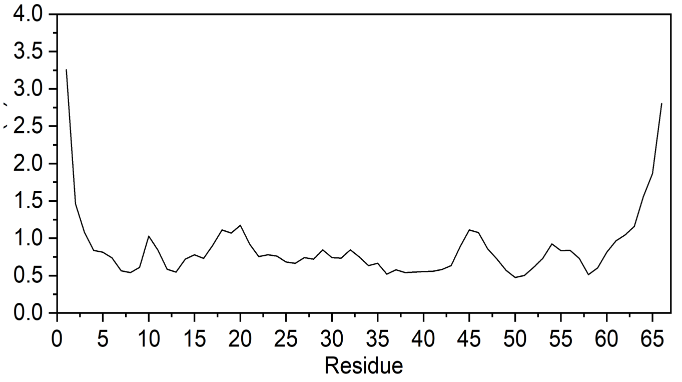

#### Supplementary Figure 6

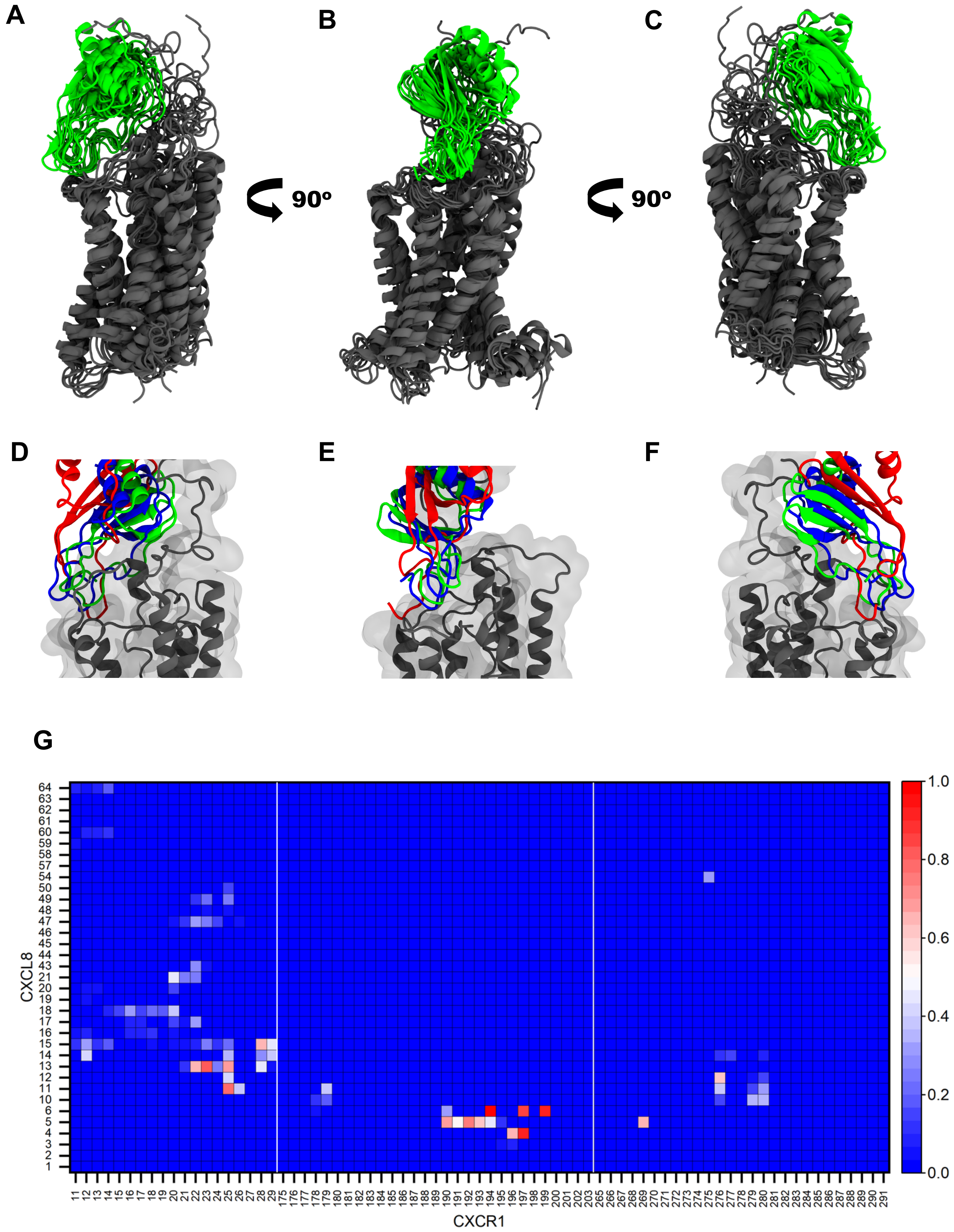

### Supplementary Figure 7

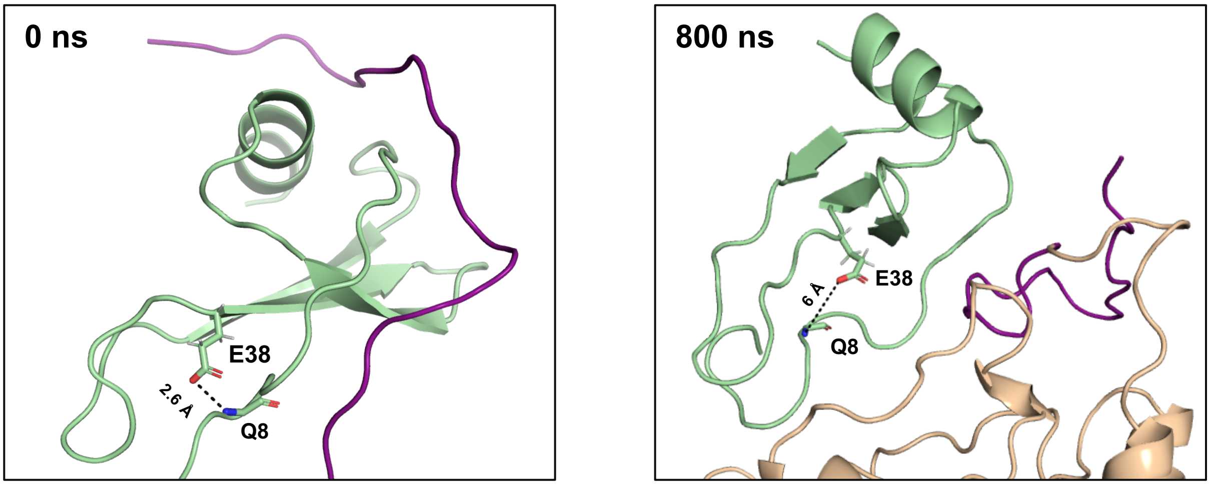

**Supplementary Figure 8**

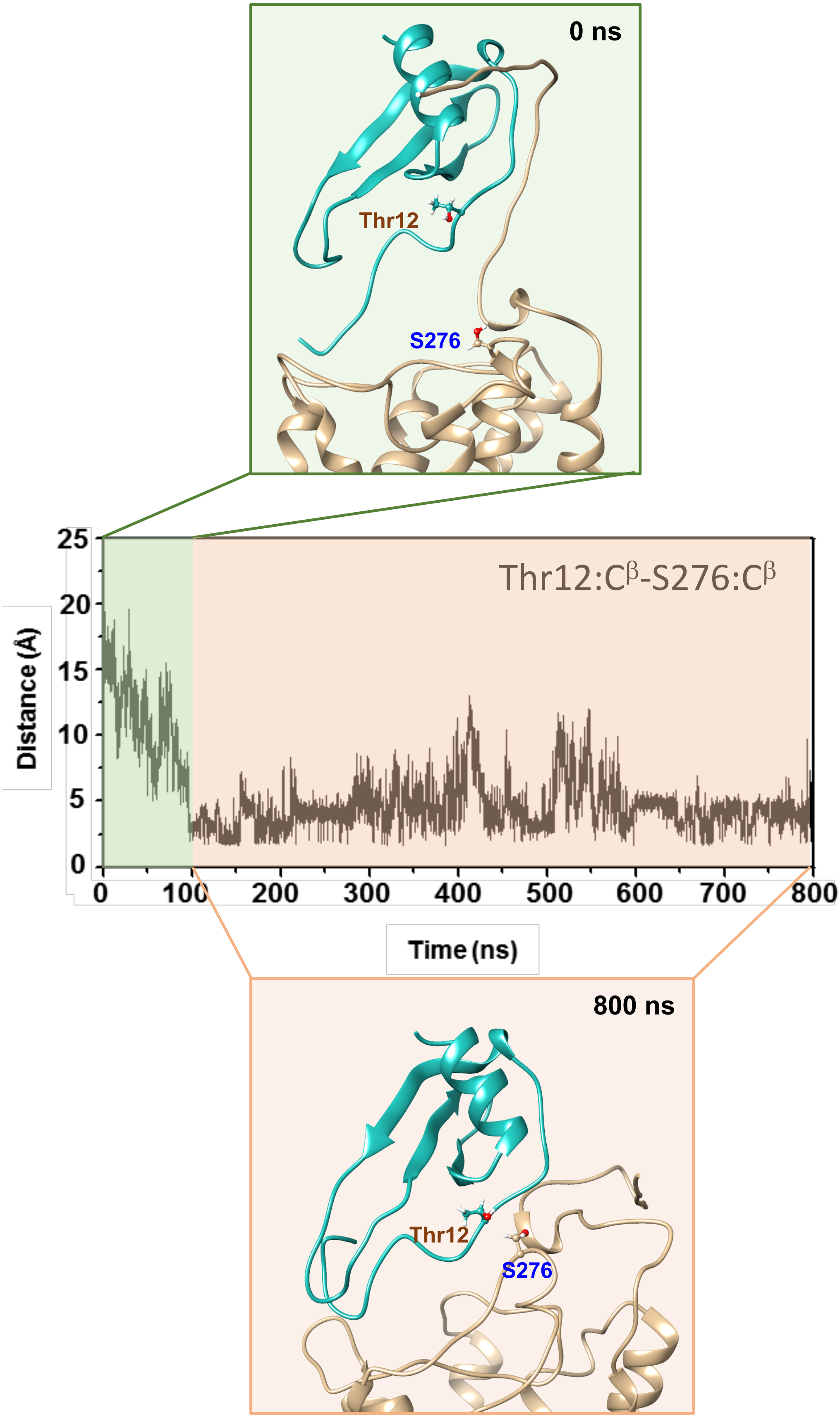

### Supplementary Figure 9

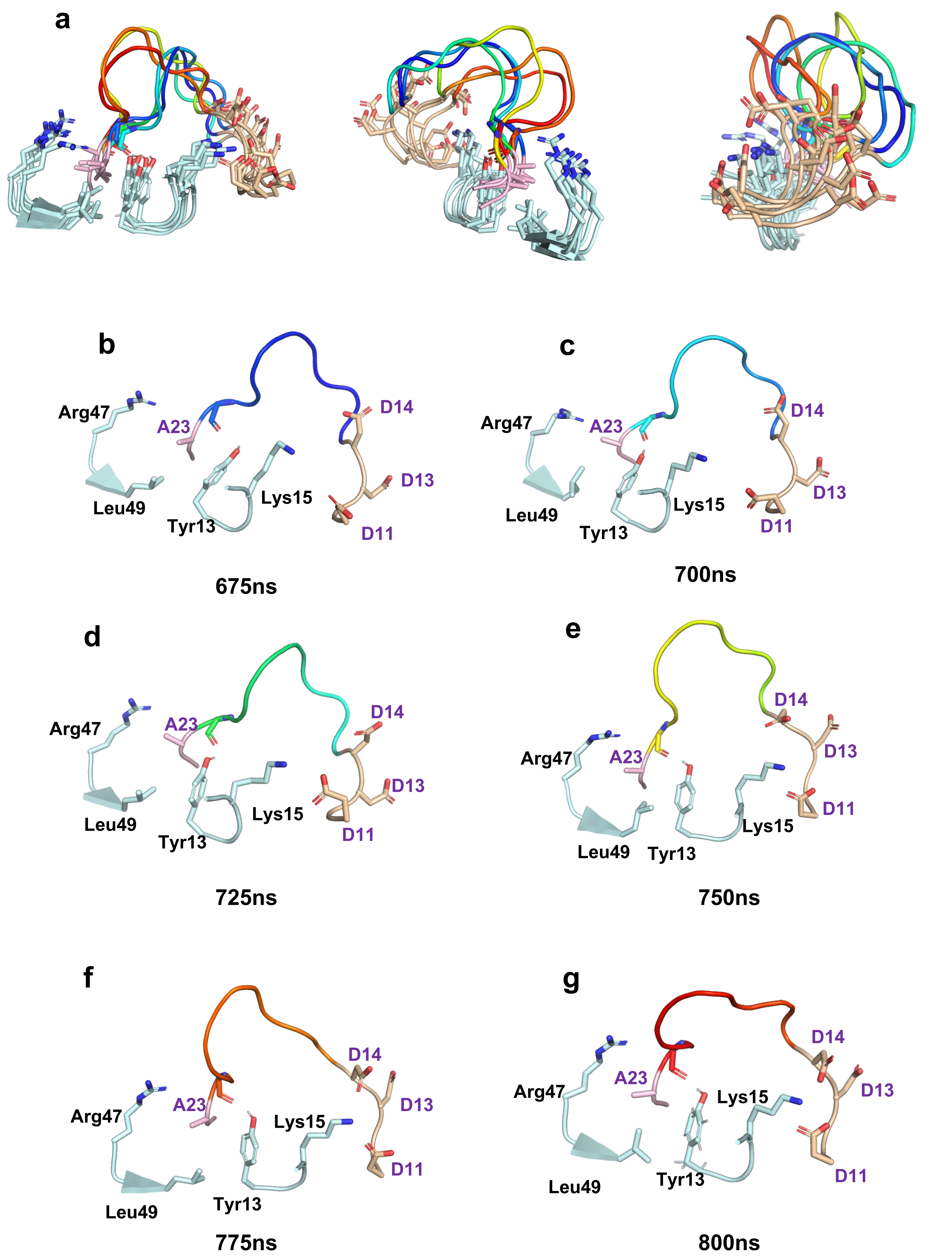

### Supplementary Figure 10

```
CXCL8          SAKELRCQCIKTYSKPFHPKFIKELRVIESGPHCANTEIIVKLSDGRELCLDPKENWVQRVVEKFLKRAENS
CXCL1        ASVATELRCQCLQTLQ-GIHPKNIQSVNVKSPGPHCAQTEVIATLKNGRKACLNPASPIVKKIIEKMLNSDKSN
CXCL2        APLATELRCQCLQTLQ-GIHLKNIQSVKVKSPGPHCAQTEVIATLKNGQKACLNPASPMVKKIIEKMLKNGKSN
CXCL3        ASVVTELRCQCLQTLQ-GIHLKNIQSVNVRSPGPHCAQTEVIATLKNGKKACLNPASPMVQKIIEKILNKGSTN
CXCL4       EAEEDGDLQCLCVKTTS-QVRPRHITSLEVIKAGPHCPTAQLIATLKNGRKICLDLQAPLYKKIIKKLLES
CXCL5    AGPAAAVLRELRCVCLQTTQ-GVHPKMISNLQVFAIGPQCSKVEVVASLKNGKEICLDPEAPFLKKVIQKILDGGNKEN
CXCL6     GPVSAVLTELRCTCLRVTL-RVNPKTIGKLQVFPAGPQCSKVEVVASLKNGKQVCLDPEAPFLKKVIQKILDSGNKKN
CXCL7            AELRCMCIKTTS-GIHPKNIQSLEVIGKGTHCNQVEVIATLKDGRKICLDPDAPRIKKIVQKKLAGDESAD
CXCL9        TPVVRKGRCSCISTNQGTIHLQSLKDLKQFAPSPSCEKIEIIATLKNGVQTCLNPDSADVKELIKKWEKQVSQ
CXCL10       VPLSRTVRCTCISISNQPVNPRSLEKLEIIPASQFCPRVEIIATMKKGEKRCLNPESKAIKNLLKAVSKEMSKRSP
CXCL11       FPMFKRGRCLCIGPGVKAVKVADIEKASIMYPSNNCDKIEVIITLKEKGQRCLNPKSKQARLIIKKVERKNF
CXCL13     VLEVYYTSLRCRCVQESSVFIPRRFIDRIQILPRGNGCPRKEIIVWKKNKSIVCVDPQAEWIQRMMEVLRKRSSSTLPVPVF
CXCL15      RPCDTQELRCLCIQEHSEFIPLKLIKNIMVIFETIYCNRKEVIAVPKNGSMICLDPDAPWVKATVGPITNRFLPEDLKQKEF
CCL1       SKSMQVPFSRC-CFSFAEQEIPLRAILCYR-NTSS-ICSNEGLIFKLKRGKEACALDTVGWVQRHRKMLRHCPSKRK
CCL2       QPDAINAPVTC-CYNFTNRKISVQRLASYRRITSS-KCPKEAVIFKTIVAKEICADPKQKWVQDSMDHLDKQTQTQTPKT
CCL3       ASLAADTPTAC-CFSYTSRQIPQNFIADY-FETSS-QCSKPGVIFLTKRSRQVCADPSEEWVQKYVSDLELSA
CCL4       APMGSDPPTAC-CFSYTARKLPRNFVVDY-YETSS-LCSQPAVVFQTKRSKQVCADPSESWVQEYVYDLELN
CCL5        SPYSSDTTPC-CFAYIARPLPRAHIKEY-FYTSG-KCSNPAVVFVTRKNRQVCANPEKKWVREYINSLEMS
CCL7       QPVGINTSTTC-CYRFINKKIPKQRLESYRRTTSS-HCPREAVIFKTKLDKEICADPTQKWVQDFMKHLDKKTQ--TPKL
CCL8       QPDSVSIPITC-CFNVINRKIPIQRLESYTRITNI-QCPKEAVIFKTKRGKEVCADPKERWVRDSMKHLDQIFQNLKP
CCL11        GPASVPTTC-CFNLANRKIPLQRLESYRRITSG-KCPQKAVIFKTKLAKDICADPKKKWVQDSMKYLDQSSPTPKP
CCL13      QPDALNVPSTC-CFTFSSKKISLQRLKSY-VITTS-RCPQKAVIFRTKLGKEICADPKEKWVQNYMKHLGRKAHTLKT
CCL14   TESSSRGPYHPSEC-CFTYTTYKIPRQRIMDY-YETNS-QCSKPGIVFITKRGHSVCTNPSDKWVQDYIKDMKEN
CCL17       ARGTNVGREC-CLEYFKGAIPLRKLKTW-YQTSE-DCSRDAIVFVTVQGRAICSDPNNKRVKNAVKYLQSLERS
CCL19         GTNDAEDC-CLSVTQKPIPGYIVRNFHYLLIKDGCRVPAVVFTTLRGRQLCAPPDQPWVERIIQRLQRTSAKMKRRSS
CCL20           ASNFDC-CLGYTDRILHPKFIVGFTRQLANEGCDINAIIFHTKKKLSVCANPKQTWVKYIVRLLSKKVKNM
CCL23      LDRFHATSADC-CISYTPRSIPCSLLESY-FETNS-ECSKPGVIFLTKKGRRFCANPSDKQVQVCMRMLKLDTRIKKTRK
CCL24          VVIPSPC-CMFFVSKRIPENRVVSYQLSSRS-TCLKAGVIFTTKKGQQSCGDPKQEWVQRYMKNLDAKQKKASPRARAVA
```

#### Supplementary Figure 11

| | |
|---|---|
| CXCR1 | MSNITDPQMWDFDDLNFTGMPPADEDYSPCML |
| CXCR2 | MEDFNMESDSFEDFWKGEDLSNYSYSSTLPPFLLDAAPCEP |
| CXCR3 | MVLEVSDHQVLNDAEVAALLENFSSSYDYGENESDSCCT |
| CXCR4 | MEGISSIPLPLLQIYTSDNYTEEMGSGDYDSMKEPCFR |
| CXCR5 | MNYPLTLEMDLENLEDLFWELDRLDNYNDTSLVENHLCPA |
| CXCR6 | MAEHDYHEDYGFSSFNDSSQEEHQDFLQFSKVFLPCMY |
| CXCR7 | MDLHLFDYSEPGNFSDISWPCNSSDCIV |
| CCR1 | METPNTTEDYDTTTEFDYGDATPCQK |
| CCR2 | MLSTSRSRFIRNTNESGEEVTTFFDYDYGAPCHK |
| CCR3 | MTTSLDTVETFGTTSYYDDVGLLCEK |
| CCR4 | MNPTDIADTTLDESIYSNYYLYESIPKPCTK |
| CCR5 | MDYQVSSPIYDINYYTSEPCQK |
| CCR6 | MSGESMNFSDVFDSSEDYFVSVNTSYYSVDSEMLLCSL |
| CCR7 | MKSVLVVALLVIFQVCLCQDEVTDDYIGDNTTVDYTLFESLCSK |
| CCR8 | MDYTLDLSVTTVTDYYYPDIFSSPCDA |
| CCR9 | MTPTDFTSPIPNMADDYGSESTSSMEDYVNFNFTDFYCEK |
| CCR10 | MGTEATEQVSWGHYSGDEEDAYSAEPLPELCYK |

### Supplementary Figure 12

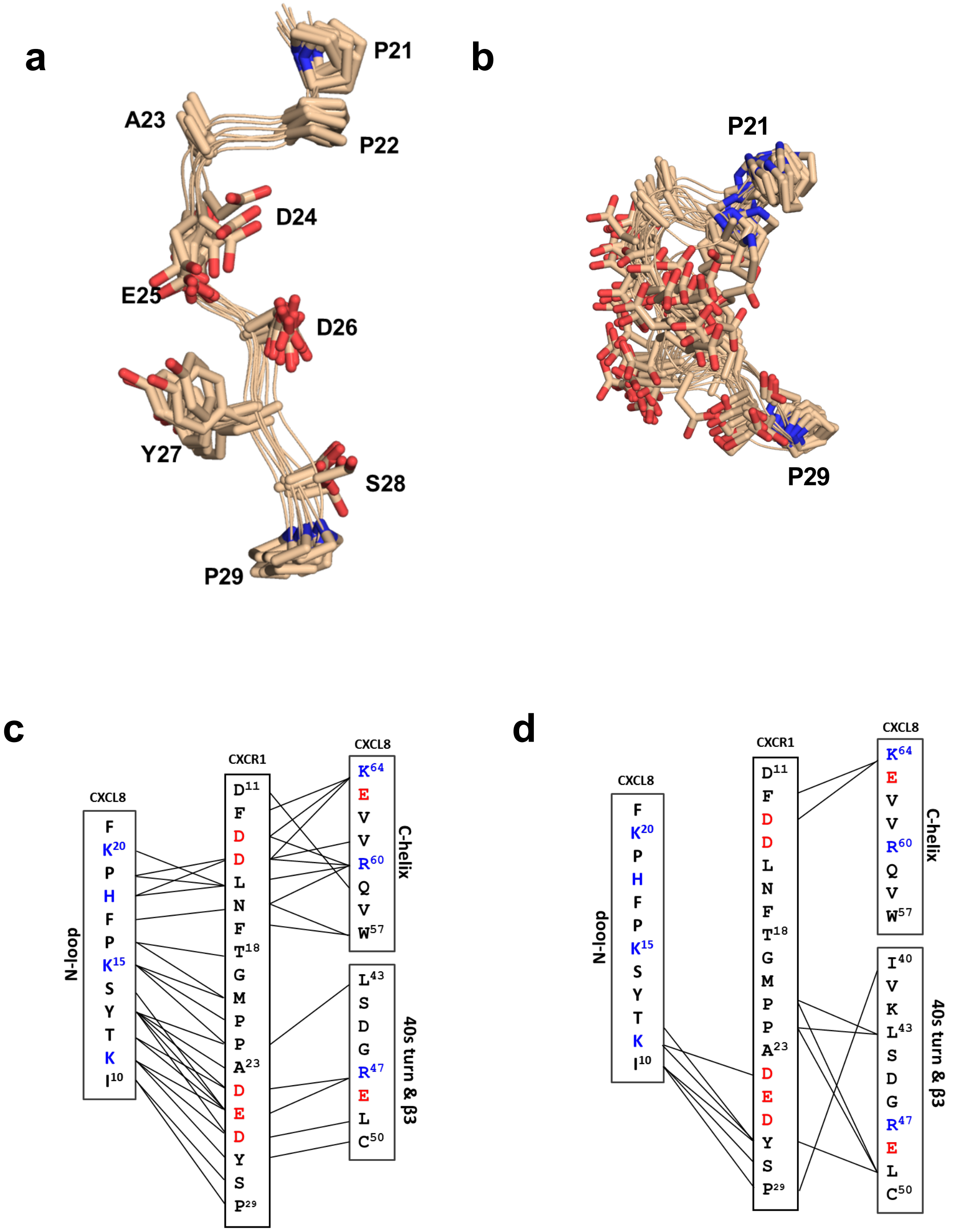

**Supplementary Table 1**. NMR Structural Statistics and r.m.s. differences for 20 calculated CXCL8-CXCR1 N-domain complex structures

| | |
|---|---|
| Energies (kcal $mol^{-1}$) | |
| NOE[a] | 3.81 ± 0.21 |
| Dihedral[a] | 0.03 ± 0.01 |
| Bonds | 3.92 ± 0.06 |
| van der Waals | 2.19 ± 0.40 |
| Deviations from idealized geometry[b] | |
| Bonds (Å) | 0.0012 ± 0.0003 |
| Angles (˚) | 0.512 ± 0.002 |
| Improper (˚) | 0.125 ± 0.001 |
| Atomic r.m.s. difference (Å)[c] | |
| **CXCL8** | |
| Backbone atoms (11-66) | 0.34 ± 0.04 Å |
| Heavy atoms (11-66) | 0.80 ± 0.06 Å |
| Regular secondary structure elements | 0.26 ± 0.02 Å |
| **CXCR1 N-domain** | |
| Backbone atoms (11-29) | 0.63 ± 0.04 Å |
| Heavy atoms (11-29) | 1.31 ± 0.06 Å |

[a]Values for NOE and torsion angles were calculated from a square well potential with a force constant of 50 kcal $mol^{-1}$ $Å^2$ and 200 kcal $mol^{-1}rad^{-2}$, respectively.
[b]Deviation for bonds, angles, and improper from ideal values based on perfect stereochemistry
[c]r.m.s. differences of the 20 final structures superimposed on the average structures

**Supplementary Table-II**: List of intermolecular NOEs

| CXCL8 | CXCR1 |
|---|---|
| ILE 10 HA | TYR 27 HB# |
| ILE 10 HG# | SER 28 HB# |
| ILE 10 HG# | PRO 29 HD# |
| ILE 10 HG# | PRO 29 HB# |
| LYS 11 HD# | TYR 27 HN |
| LYS 11 HD# | TYR 27 HD# |
| LYS 11 HB# | ASP 26 HB# |
| LYS 11 HD# | ASP 26 HB# |
| LYS 11 HB# | GLU 25 HB# |
| LYS 11 HD# | GLU 25 HB# |
| THR 12 HB | ASP 26 HB# |
| THR 12 HB | GLU 25 HB# |
| TYR 13 HB# | ASP 26 HB# |
| TYR 13 HB# | ASP 24 HG# |
| TYR 13 HB# | ASP 24 HA |
| TYR 13 HB# | GLU 25 HB# |
| TYR 13 HB# | PRO 22 HD# |
| TYR 13 HD# | ASP 26 HB# |
| TYR 13 HD# | GLU 25 HB# |
| TYR 13 HE# | ALA 23 HB# |
| SER 14 HB# | GLU 25 HB# |
| LYS 15 HB# | MET 20 HG# |
| LYS 15 HG# | MET 20 HB# |
| LYS 15 HG# | PRO 21 HB# |
| LYS 15 HG# | PRO 22 HD# |
| LYS 15 HD# | PRO 21 HB# |
| LYS 15 HD# | PRO 21 HD# |
| LYS 15 HD# | PRO 22 HB# |
| LYS 15 HE# | MET 20 HA |
| PRO 16 HB# | MET 20 HB |
| PRO 16 HD# | MET 20 HB |
| PRO 16 HD# | THR 18 HB |
| PHE 17 HB# | ASN 16 HB# |
| PHE 17 HB# | ASN 16 HA |
| PHE 17 HD# | PHE 17 HB# |
| HIS 18 HA | ASP 14 HB# |
| HIS 18 HB# | ASP 14 HB# |
| HIS 18 HB# | LEU 15 HG |
| HIS 18 HB# | LEU 15 HB# |

| | |
|---|---|
| HIS 18 HD# | ASP 14 HB# |
| HIS 18 HD# | LEU 15 HG |
| HIS 18 HD# | LEU 15 HB# |
| PRO 19 HA | ASP 14 HB# |
| PRO 19 HB# | ASP 14 HB# |
| PRO 19 HB# | LEU 15 HG |
| PRO 19 HG | LEU 15 HG |
| PRO 19 HD# | ASP 14 HB# |
| PRO 19 HD# | LEU 15 HB# |
| LYS 20 HG# | LEU 15 HG |
| LYS 20 HG# | LEU 15 HB# |
| LYS 20 HD# | LEU 15 HA |
| LEU 43 HB# | PRO 22 HA |
| LEU 43 HB# | PRO 22 HG# |
| LEU 43 HG | PRO 22 HG# |
| LEU 43 HG | PRO 21 HB# |
| LEU 43 HD# | PRO 21 HG# |
| ARG 47 HB# | ASP 24 HB# |
| ARG 47 HB# | GLU 25 HB# |
| ARG 47 HG# | GLU 25 HG# |
| ARG 47 HG# | ASP 24 HB# |
| ARG 47 HD# | ASP 24 HB# |
| ARG 47 HD# | GLU 25 HB# |
| LEU 49 HB# | ASP 26 HA |
| LEU 49 HB# | GLU 25 HB# |
| LEU 49 HG | ASP 26 HB# |
| LEU 49 HG | GLU 25 HB# |
| TRP 57 HB# | LEU 15 HB |
| TRP 57 HB# | ASN 16 HB# |
| TRP 57 HD1 | ASN 16 HA |
| TRP 57 HE1 | ASN 16 HB# |
| ARG 60 HG# | ASP 14 HB# |
| ARG 60 HG# | ASP 13 HB# |
| ARG 60 HD# | ASP 14 HB# |
| ARG 60 HD# | LEU 15 HB# |
| ARG 60 HD# | LEU 15 HD# |
| ARG 60 HD# | ASN 16 HB# |
| ARG 60 HD# | ASP 13 HA |
| ARG 60 HE# | ASP 14 HB |
| VAL 61 HB | ASP 14 HA |
| VAL 61 HB | ASP 14 HB# |
| LYS 64 HG# | ASP 13 HB# |

| | |
|---|---|
| LYS 64 HG# | ASP 13 HB# |
| LYS 64 HG# | ASP 13 HB# |
| LYS 64 HG# | ASP 14 HB# |
| LYS 64 HD# | ASP 14 HB# |
| LYS 64 HE# | ASP 14 HB# |
| LYS 64 HE# | PHE 12 HB# |
